# Epigenetic regulation of translation repression in ferroptosis, and a role of Alternative splicing and tRNA methylation

**DOI:** 10.1101/2021.11.10.468143

**Authors:** Sherif Rashad, Daisuke Saigusa, Yuan Zhou, Liyin Zhang, Teiji Tominaga, Kuniyasu Niizuma

## Abstract

Ferroptosis is a non-apoptotic cell death mechanism characterized by the production of lipid peroxides. Ferroptosis plays important roles in many diseases such as cancer and neurodegenerative diseases. While many effectors in the ferroptosis pathway have been mapped, its epigenetic and epitranscriptional regulatory processes are not yet fully understood. Ferroptosis can be induced via system xCT inhibition (Class I) or GPX4 inhibition (Class II). Previous works have revealed important differences in cellular response to Class I and Class II ferroptosis inducers. Importantly, blocking mRNA transcription or translation appears to protect cells against Class I ferroptosis inducing agents but not Class II. Understanding these subtle differences is important in understanding ferroptosis as well as in developing therapeutics based on ferroptosis for various diseases. In this work, we examined the impact of blocking transcription (via Actinomycin D) or translation (via Cycloheximide) on Erastin (Class I) or RSL3 (Class II) induced ferroptosis. Blocking transcription or translation protected cells against Erastin but was detrimental against RSL3. Cycloheximide led to increased levels of GSH alone or when co-treated with Erastin and the activation of the reverse transsulfuration pathway. RNA sequencing analysis revealed an important and unexplored role of Alternative splicing (AS) in regulating ferroptosis stress response and mRNA translation repression. Our results indicated that translation repression is protective against Erastin but detrimental against RSL3. We tested this theory in Alkbh1 overexpressing glioma cells. Alkbh1 demethylates tRNA and represses translation and is associated with worse outcome in glioma patients. Our results showed that Alkbh1 overexpression protected glioma cells against Erastin but was detrimental against RSL3.

## Introduction

Ferroptosis is a caspase-independent cell death mechanism that was discovered nearly a decade ago^1^. Since its reporting, ferroptosis-mediated cell death has been implicated in an exponentially increasing number of disease mechanisms, including cancer, neurodegeneration, and ischemic injuries^2–4^. Large strides have been made in understanding the genetic and metabolic regulators of ferroptosis, with the advent of CRISPR and chemical screening as well as other high throughput approaches^5–7^. These approaches succeeded in mapping various effectors in the ferroptosis pathway and identifying different classes of ferroptosis inducing agents^8^.

Despite these important advances in our understanding of ferroptosis, we are still scratching the surface when it comes to fully grasping the nuances of this interesting process. While a large body of work has focused on the metabolic regulators of ferroptosis^9^, the epigenetic and epitranscriptional regulators of ferroptosis are yet to be fully explored. For example, it is known that RNA and tRNA modifications play important roles in regulating cellular sensitivity to oxidative stresses^10,11^, however, these mechanisms are not fully explored in the context of ferroptosis. Furthermore, recent studies revealed a link between ferroptosis and the integrated stress response^12^ and mitochondrial stress^13^. Both mechanisms are known to be heavily regulated at the level of epigenome and epitranscriptome. Important differences also exist in cellular metabolic response to ferroptosis induced by different classes of ferroptosis inducers^14^. Interestingly, blocking translation via cycloheximide (CHX) appears to have different effects on ferroptosis induction via system xCT blockade or via GPX4 inhibition^5,15^. Protein translation repression is an integral part of cellular response to oxidative stress, and it is mediated by complex and intertwining transcriptional, epigenetic, and epitranscriptional mechanisms. Understanding how these mechanisms contribute to cellular response to ferroptosis would be paramount in our understand of ferroptosis regulation.

In this work we examined the impact of transcription and translation blockade on cellular response to ferroptosis. We identified a strong Alternative splicing (AS) program being activated during ferroptosis prior to transcriptional reprogramming, yet this program was different between system xCT inhibition and GPX4 inhibition induced ferroptosis. Finally, we identify a role of tRNA modifications via the tRNA demethylase Alkbh1 in dictating glioma cells sensitivity to ferroptosis.

## Results

### Effect of translation or transcription inhibition on ferroptosis

To interrogate the effect of transcription or translation inhibition on ferroptosis, we started by treating B35 neuroblastoma cells with the System xCT inhibitor Erastin (Class I ferroptosis inducing agent) or with the GPX4 inhibitor RSL3 (Class II ferroptosis inducing agent). We cotreated the cells with the transcription inhibitor Actinomycin D (ActD) or the translation inhibitor cycloheximide (CHX) or with vehicle (DMSO). We adjusted the drug concentrations or Erastin and RSL3 to reach approximately 40~50% cell viability on WST-8 assay after 8 hours of stress [Figure 1a]. As previously observed^5,16^, blocking translation or transcription both protected cells treated with Erastin. On the other hand, ActD had no effect on RSL3 treated cells while CHX greatly enhanced the cell death [Figure 1a & b]. Next, we examined the presence of lipid peroxides with Bodipy C11 staining and the effect of ActD and CHX on their generation. Lipid peroxides were robustly expressed 4 hours after RSL3. While in the Erastin treatment group, they were observed 6 hours after treatment. The differences in the time points of observation are due to the fact that while the endpoint of lipid peroxidation exist in both groups, there are mechanistic differences in their generation between the 2 drugs^14^. In the RSL3 group, ActD treatment resulted in comparable lipid peroxide generation with the RSL3 only group. CHX, on the other hand, resulted in extensive cell death that interfered with the observation of lipid peroxides at the chosen timepoint [Figure 1c]. Ferrostatin-1 (Ferr1; ferroptosis inhibitor), greatly mitigated the impact of RSL3 and RSL3+ActD on the cells. It also reduced the level of cell death and detachment observed when the cells were treated with RSL3+CHX, however, there was still extensive lipid peroxidation observable [Figure 1d]. In the Erastin treatment group, Lipid peroxides were observed at 6 hours [not shown] and 8 hours along with extensive cell death [Figure 1e]. Co-treatment with ActD, CHX, or Ferr1 abrogated the cell death and lipid peroxidation [Figure 1e]. These results are in-line with previous observations^5,16^ and indicate that transcription or translation repression, whether innate or induced, can be advantageous when ferroptosis is induced by Class I FIN (Ferroptosis inducers) which act on system xCT, but detrimental with Class II FIN which act on GPX4.

**Figure 1:**
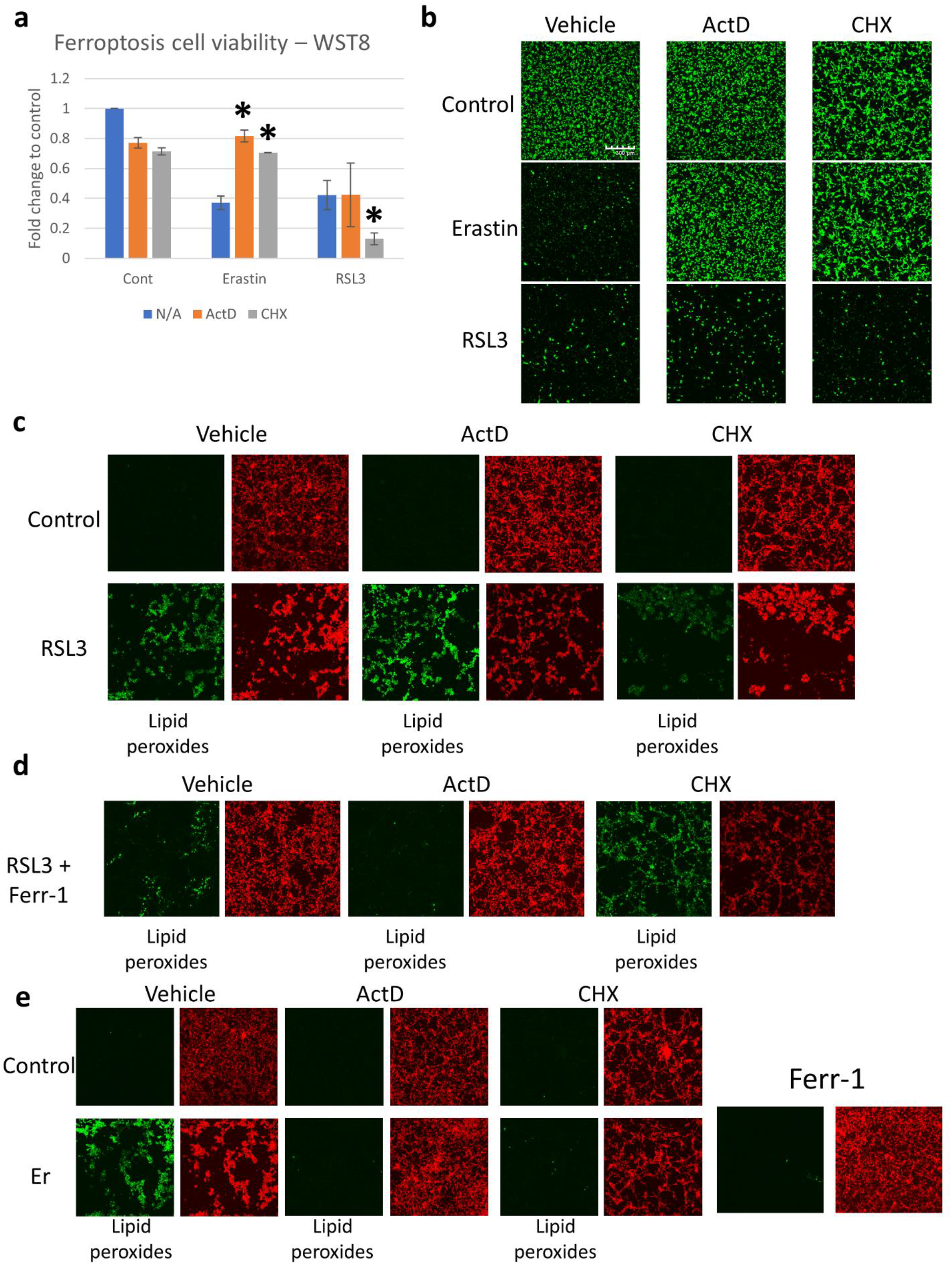
Blocking transcription or translation rescues cells from Erastin but not RSL3 induced ferroptosis. B35 cells were incubated with Erastin (1μM) or RSL3 (0.1μM) in addition to Actinomycin D (ActD; 5μg/ml) or Cycloheximide (CHX; 100μg/ml). a: WST-8 assay after 8 hours of incubation showing recovery of cell viability when cells were co-treated by ActD or CHX and Erastin, while CHX worsened cell viability when co-treated with RSL3 (Experiment repeated thrice, 6 replicates each). Asterisk = statistically significant (*p* < 0.05) versus vehicle (N/A) group. b: Calcein green live cell imaging showing rescue of cell viability in the Erastin group with ActD or CHX co-treatment and the absence of effect in the RSL3 group (experiment repeated thrice with 2 replicates each). Scale bar = 300μm c: Bodipy C11 live cell imaging 4 hours after exposure to RSL3 with ActD or CHX co-treatment showing the generation of lipid peroxides in all groups. RSL3 + CHX led to extensive cell death. Green = lipid peroxides. Red = unoxidized lipids. d: Ferrostatin-1 (Ferr-1) reduced lipid peroxidation after RSL3 and RSL3 + ActD treatment. Ferr-1 rescued cell viability to some extent in the RSL3 + CHX group, but Lipid peroxides were still evident. e: Bodipy C11 staining in the Erastin group showing that ActD, CHX, and Ferr-1 all blocked lipid peroxides formation when co-treated with Erastin (Er). Bodipy experiments were repeated three times with two replicates per experiment.

### Paradoxical effect of CHX on GSH when combined with Erastin or RSL3

Ferroptosis is centered around the detoxifying activity of GPX4 along with reduced glutathione (GSH)^17^. In order to evaluate the impact of transcription or translation inhibition on GSH, and other relevant metabolites, we performed liquid chromatography mass spectrometry (UHPLC-MS/MS), using the same groups shown in figure 1a, after 8 hours of ferroptosis stress exposure. ActD treatment, without RSL3 or Erastin, reduced GSH levels. On the other hand, CHX slightly increased GSH levels [Figure 2a]. While this might appear counter intuitive, translation repression during stress was shown to shift the mRNA translation machinery towards the generation of stress proteins, and importantly GSH, via changes in transfer RNA (tRNA) transcripts expression levels and consequently mRNA codon usage and translation rates^18^. Whether CHX acts via a mechanism simulating this effect remains to be examined. Erastin treatment caused GSH depletion, in line with its known mechanism of action^1^. ActD paradoxically increased the levels of GSH when cells were co-treated with Erastin, despite its GSH lowering effect in the absence of Erastin or RSL3, and CHX co-treatment greatly recovered the GSH levels [Figure 2a]. RSL3 co-treatment with ActD or CHX lowered GSH levels, which appears paradoxical to the activity of CHX with Erastin or by itself. This might be a consequence of the extensive cell death caused by the co-treatment with RSL3 and CHX [Figure 2a]. Oxidized glutathione (GSSG) was not detected after Erastin treatment, which can correlate with GSH depletion and the unavailability of GSH pools to interact with lipid peroxides [Figure 2b]. This can be extrapolated from the detection of GSSG when CHX is used with Erastin, which led to GSH generation and therefore the availability of GSH to detoxify lipid peroxides.

**Figure 2:**
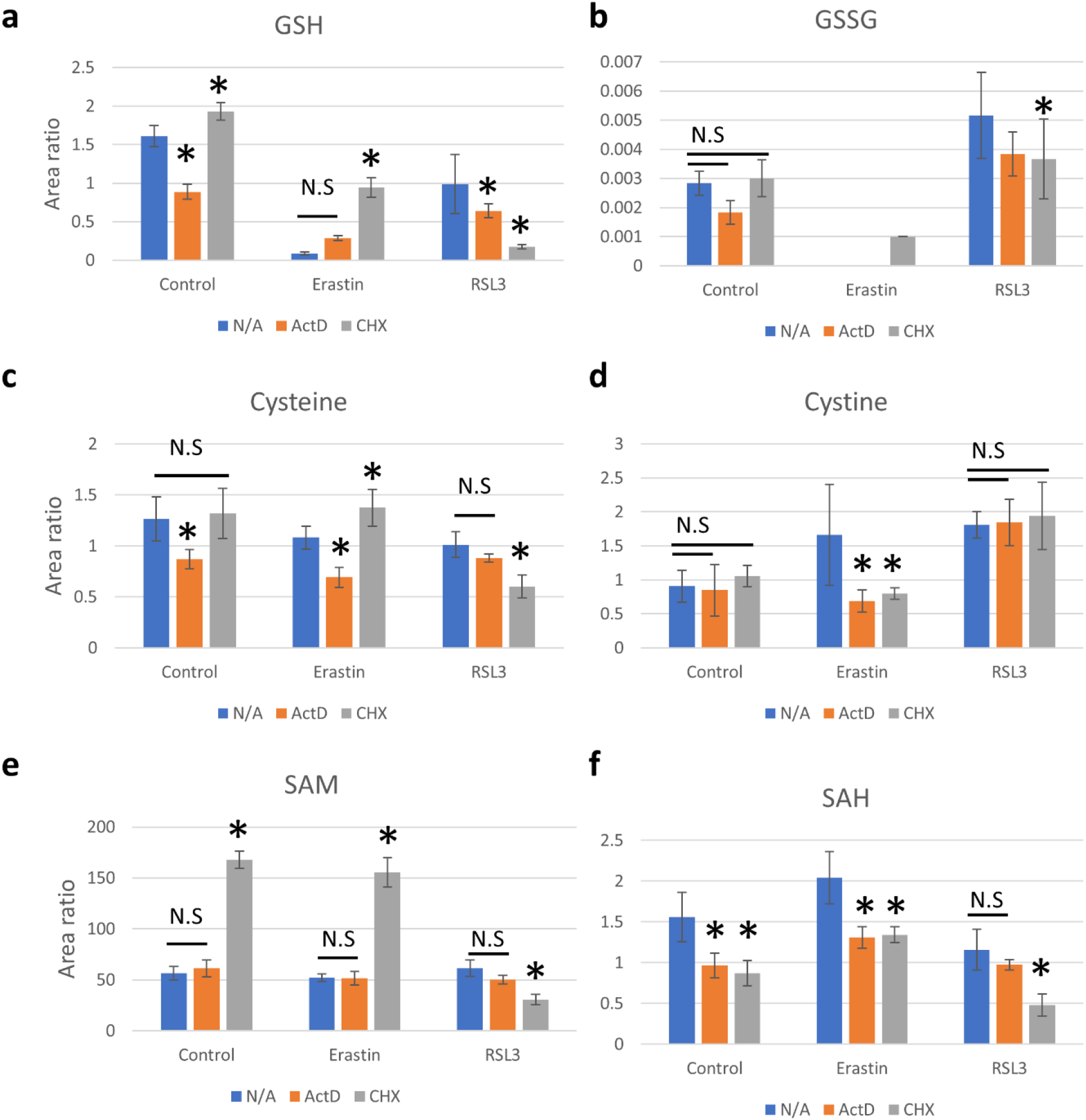
Cycloheximide rescues Glutathione depletion in Erastin and elevates SAM. Mass spectrometry analysis showing the effect of ActD and CHX co-treatment with Erastin or RSL3 on the levels of a: reduced glutathione (GSH), b: oxidized glutathione (GSSG), c: Cysteine, d: Cystine, e: S-adenosyl methionine (SAM), and f: S-adenosyl homocysteine (SAH). Data is presented as Area ratio. Equal cell numbers were used for all experiments. Mass spectrometry was conducted with three replicates per group. Asterisk = statistically significant (*p* < 0.05) versus vehicle (N/A) treated group. See supplementary figure 1.

Several other related metabolites also showed similar changes that can be correlated with cell viability and lipid detoxification [Figure 2c-f, Supplementary figure 1]. Surprisingly, CHX caused an increase in cysteine (Cys) levels, the precursor for GSH synthesis^8^, when combined with Erastin, without and apparent increase in intracellular cystine, which is imported via the system xCT in exchange for glutamate^8^ [Figure 2c]. ActD, on the contrary, reduced Cys levels alone and with Erastin co-treatment. Which appears contradictory to the need for Cys for GSH synthesis and ferroptosis protection. It is notable that the levels of Cys did not change from control when RSL3 or Erastin were used alone. Looking at cystine levels, it is clear that both ActD and CHX reduced cystine when combined with Erastin [Figure 2d]. The reasons for these changes are yet to be understood.

S-adenosylmethionine (SAM), the main methyl donor in the cell^19^, was extensively upregulated with CHX treatment or co-treatment with Erastin [Figure 2e]. SAM is known to play important role in oxidative stress^20^, however, its role in ferroptosis is yet to be established. Furthermore, the role of SAM in the protective protein translation repression response to stress, which might be mimicked by CHX in this case, is not known. On the contrary, RSL3 co-treatment with CHX reduced SAM levels, which might contribute to the increased cell death observed, or it might be a byproduct of that cell death. S-adenosyl homocysteine (SAH), another metabolite which is known to play important role in oxidative stress was also affected by ActD and CHX treatment. Both drugs decreased the levels of SAH alone or with Erastin co-treatment. While with RSL3, only CHX reduced SAH levels significantly [Figure 2f]. SAH was shown to correlate and promote oxidative stress^21^, which can explain its increase with Erastin treatment. ActD and CHX offered protection against Erastin induced ferroptosis might act via reducing its levels along with other mechanisms. It’s important to note that SAH levels did not increase with RSL3, which might indicate that SAH does not play an important role in RSL3 mediated ferroptosis, and its reduction with CHX co-treatment might be an effect rather than a cause.

The upregulation of GSH, Cysteine, and SAM in the CHX group might indicate the activation of the reverse transsulfuration pathway, which was shown recently to protect against ferroptosis^22–24^. Other metabolites also hinted to the activation of this pathway. Methionine was upregulated with CHX treatment, although its levels were comparable to the vehicle group after Erastin and CHX co-treatment [Supplementary figure 1a]. Cystathionine, the precursor for Cysteine was also upregulated when CHX was co-treated with Erastin [Supplementary figure 1b]. Homocysteine also showed the same pattern as with Methionine [Supplementary figure 1c].

Overall, these results indicate that CHX and protein translation repression protects cells from Erastin induced lipid peroxidation via a mechanism that mimics cellular response to stress via translation repression, leading to generation of GSH. Both ActD and CHX might also protect the cells via reducing SAH and CHX might be acting to increase SAM levels, offering an antioxidant advantage to the cells. The transsulfuration pathway appears to play a role in CHX mediated protection only, and not in ActD. These mechanisms do not appear to be functioning in RSL3 mediated ferroptosis.

### Transcriptional and epigenetic response to Erastin and RSL3

Our data indicates that transcription and translation play important roles in ferroptosis. Guided by this observation, we shifted our focus to the transcriptome in order to further understand the cellular response to ferroptosis. We chose to perform mRNA sequencing 4 hours after exposure to the drugs, due to the fact that at 8 hours, we observed extensive cell death especially in the RSL3+CHX group as evident from the cell viability analysis and from the reduction of many of the metabolites analyzed, which can be detrimental to the quality of RNA sequencing.

We first started by analyzing the differentially expressed genes (DEG) after RSL3 or Erastin treatment. Erastin exposure for 4 hours did not have impact on differentially expressed genes as compared to the control [Figure 3a]. We believe this is due to the earlier time point selected that precedes lipid peroxide production after Erastin treatment. RSL3, on the other hand, showed less than 100 statistically significant DEGs as compared to the control samples [Figure 3a]. Gene ontology (GO) and Gene set enrichment (GSE) and pathway analysis revealed an enrichment for stress induced transcription in the RSL3 treated cells [Figure 3b] as the top enriched term, followed by positive regulation of cell death. This can partially explain the enhanced cell death after CHX, as blocking the translation of essentially transcribed genes constituting this transcriptional response could be detrimental to the cells if these genes act as buffer to mitigate oxidative damage.

**Figure 3:**
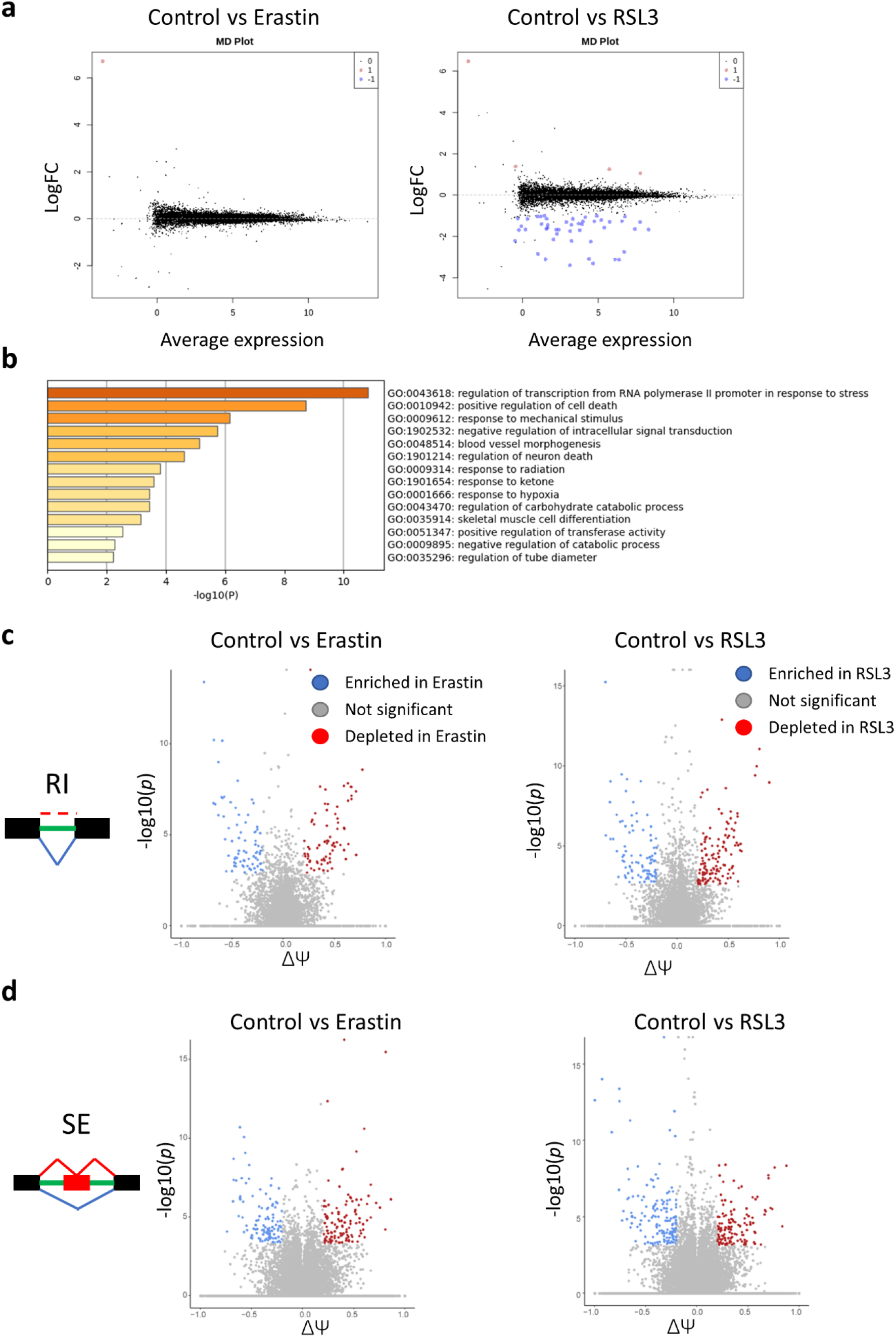
RNA-seq revealed strong activation of Alternative splicing (AS) in Ferroptosis. a: MD plots of differentially expressed genes (DEGs) 4 hours after Erastin or RSL3 exposure. No DEGS were observed in the Erastin group, while less than 100 DEGs were observed in the RSL3 group. b: Pathway analysis for the genes upregulated after RSL3 treatment. c and d: Volcano plots revealing the enrichment and depletion of hundreds of AS sites following both Erastin and RSL3 treatment. RI: retained introns. SE: skipped exons. ΔΨ: delta-psi.

We next started looking deeper into the sequencing data, as the effect on transcription was weaker than what we expected and even absent with Erastin treatment, despite a robust cell death at the end point. Oxidative stress can induce a variety of responses in the cells, and not restricted to transcription. Alternative splicing (AS) is well known mechanism that regulates cellular proteome diversity, protein translation, mRNA stability and mRNA expression as well as being a part of the oxidative stress response^25–27^. To explore the role of AS in ferroptosis response, we focused on local splice variants (LSV) analysis as our chosen method to analyze various AS types. We used the rMATs software^28^ which is capable of detecting and comparing various forms of alternative splicing: Skipped exons (SE), Retained Introns (RI), Alternative 5’ splice site (A5SS), Alternative 3’ splice site (A3SS), and mutually exclusive exons (MXE). Since each splice type would have an effect on mRNA dynamics, this classification offers an interesting way of studying AS in ferroptosis response. For example, RI can affect introduce premature stop codons in the open reading frame (ORF) leading to translation termination^27,29^, but instead of leading to mRNA degradation, mRNAs with RI are more stable^30^ and remain in the RNA pool. Thus, in this example, the changes caused by RI would not be reflected on DEG data.

AS analysis revealed hundreds of genes to be differentially alternatively spliced (D-AS) in the Erastin and RSL3 groups in each of the AS types tested as compared to control samples [Figure 3c and d, Supplementary figures 2, Supplementary tables 1 and 2]. The AS program enriched in each dataset was vastly different, as evident by the very low overlap between genes that underwent AS in each AS subtype in the Erastin and RSL3 datasets [Supplementary tables 1 and 2]. Rather than focusing on single spliced genes or event, we opted to look into the AS events as a collective, arguing that they might be regulated via a common mechanism or themselves regulating certain aspects of ferroptosis response. To do so, we performed GO analysis using genes in each AS type as input to Metascape^27^ via the multiple gene lists feature to evaluate which AS type was predominantly regulates certain pathways or processes. We analyzed the 3 gene ontology (GO) terms (GO Biological processes, GO Molecular functions, GO cellular components) for D-AS genes in the Control vs Erastin and Control vs RSL3 sets.

Looking into the number of events and overlap between spliced genes between AS events following Erastin treatment, RI, SE, and A3SS events had nearly equal number of genes. There was, however, minimal overlap between the genes present in each set [Figure 4a]. In the RSL3 treatment group, the same pattern was present [Figure 4b]. While several GO terms overlapped between various AS types in the Erastin group [Figure 4a], the overlap was much lower for the SE events in relation to other AS types in the RSL3 group [Figure 4b]. Analysis of the top enriched terms in the Erastin group [Figure 4c, supplementary figure 3] showed various terms to be enriched. The top terms were significantly enriched for protein translation with terms such as “translation”, “cytosolic ribosome”, and “translation initiation”. Moreover, RNA splicing, transcription and other epigenetic processes were also enriched. Furthermore, it appears that RI events play important role in regulating Erastin response, as most top enriched terms were enriched in the RI dataset alone or in combination with other AS types [Supplementary figure 3]. Cluster analysis of the enriched terms revealed 2 main clusters in the top enriched terms that can be divided into a “translation” related cluster and “transcription” related cluster [Supplementary figure 4a]. RI events were the major contributor to these 2 major clusters [Supplementary figure 4b]. This data puts emphasis on these processes and their related epigenetic regulatory mechanisms in the Erastin stress response as well as the significant role of RI in their regulation.

**Figure 4:**
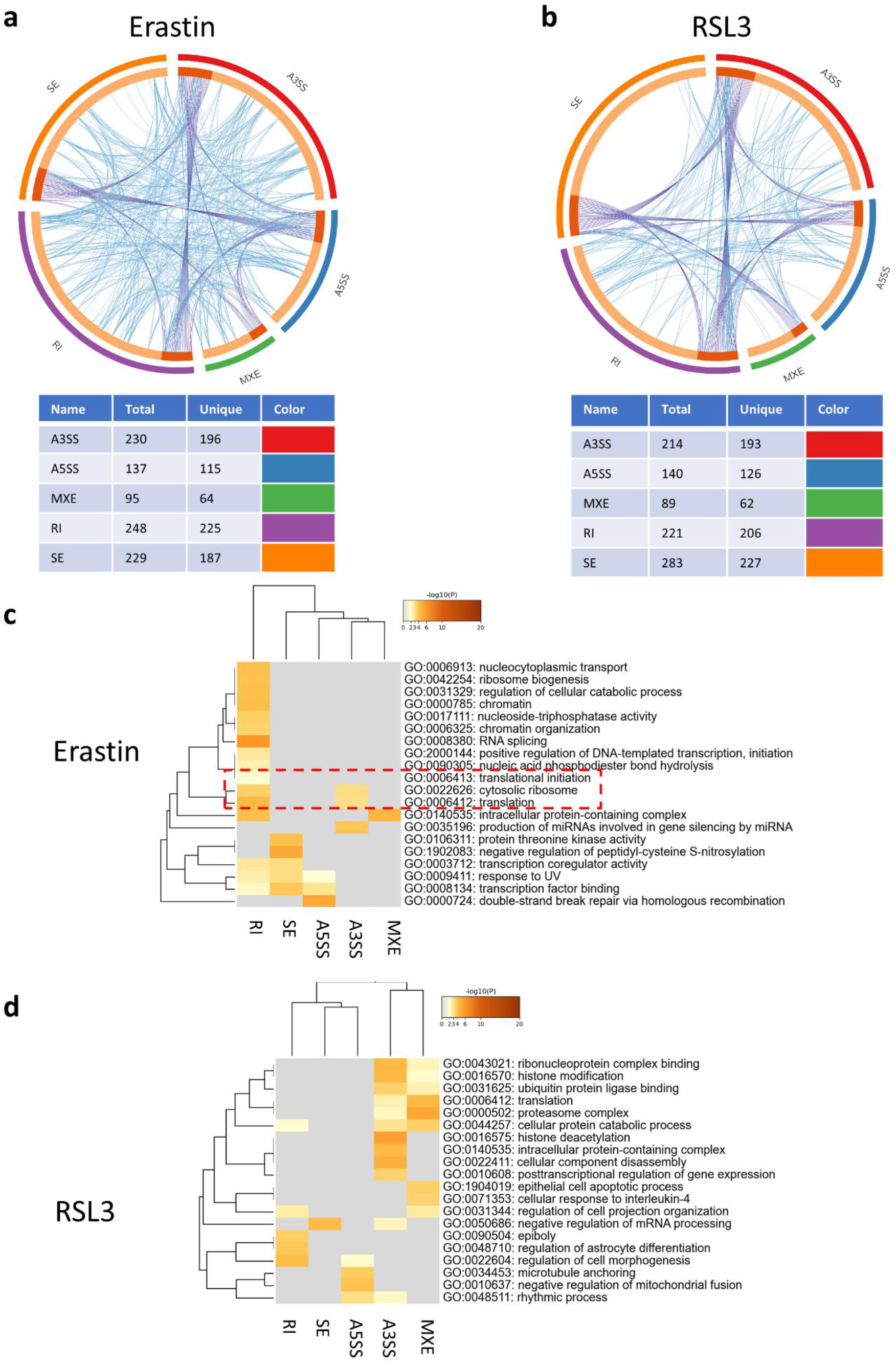
Alternative splicing events regulate various pathways impacting ferroptosis stress response. a and b: Circos plots showing the overlap between different AS events at the gene level (purple lines) or at the pathway enrichment level (blue lines) in Erastin (a) and RSL3 (b). The tables under the plots show the number of genes in each AS type and the number of unique genes belong to each AS event type. c: Gene ontology (GO) analysis for the alternatively spliced genes in the Erastin group showing a dominance of RI events in regulating multiple pathways and the presence of a node related to regulation of translation (red dotted box). d: GO analysis for the AS genes in RSL3.

In the RSL3 dataset, the top enriched terms were related to histone modifications and regulation and various protein-related processes such as proteasome activity and ubiquitin ligase binding [Figure 4d, Supplementary figure 5]. Translation relevant mechanisms although were present, were not as high scoring as in the Erastin dataset. The dominance of RI in regulating the enriched pathways observed with Erastin was not observable in the RSL3 dataset. While RI was still important, A3SS was enriching more highly scoring pathways in the RSL3 AS analysis [Supplementary figure 5]. Cluster analysis revealed several small clusters, different from the major clustering pattern observed after Erastin treatment [Supplementary figure 6].

Altogether, this analysis evidently shows that AS plays important unexplored role in cellular response to ferroptosis. This analysis also shows that AS mediated ferroptosis response is drastically different between Erastin and RSL3, with pathways regulating translation and transcription scoring high in Erastin, while pathways related to histone modifications and proteasome pathways scoring high in RSL3. Furthermore, the AS event type dominant in each group was different, with intron retention (RI) appearing to be playing important role in Erastin response.

### Influence of ActD and CHX on ferroptosis

Given that epigenetic regulation of translation or transcription appear to play important part in ferroptosis response, we shifted our attention to the impact of ActD and CHX on gene expression following ferroptosis. To do so, we first compared ActD or CHX treated groups with their ferroptosis counterparts (e.g., ActD vs RSL + ActD and so on). This analysis did not yield significant DEGs in the Erastin treatment groups with both ActD and CHX, as well as in the RSL treatment group with ActD [Supplementary figure 7]. Comparing CHX with RSL + CHX yielded 124 DEGs, however, pathway analysis was not informative to provide cell stress or ferroptosis relevant mechanisms that can explain the pattern [not shown].

The impact of ActD or CHX on Erastin or RSL3 might be related to the mechanism by which each drug impact cellular transcription and translation, in return blocking certain pathways that can impact ferroptosis stress response. To that extend we started looking into the effect that CHX would have on the transcriptional landscape and how that could be used to explain the observed effect on cell viability and lipid peroxidation. CHX acts by “freezing” ribosomes on mRNAs, thus blocking translation^31^. It is frequently used for translation inhibition during ribosome profiling experiments^32^. It was also shown to stimulate ribosome biogenesis in budding yeast at the level of transcription^31^. However, the transcriptional impact of CHX on eukaryotes is not known, let alone the impact of long exposure to CHX as in the case of our experiment. Analysis of DEGs revealed nearly 3000 genes to be differentially expressed between the control group and CHX only treatment group (1075 downregulated and 1906 upregulated in CHX) [Figure 5a]. GO enrichment analysis revealed hundreds of terms to be enriched. Looking into the top enriched terms, several processes related to DNA and chromatin activity were observed [Figure 5b]. The clustering pattern of the pathway enrichment showed a major cluster related to gene silencing, nucleosome regulation, and chromatin organization [Figure 5c, gene expression regulation cluster]. This cluster was exclusively upregulated in CHX [Supplementary figure 8]. From the analysis of AS events in the Erastin group, it is clear that RI plays important role in the expected effect. RI can delay mRNA export from the nucleus and greatly affect the translation of transcript, effectively silencing the mRNAs^27,33^. Thus is Erastin, we expect to observe a transcription and translation repressive response guided by RI. Whether the activation of various pathways that regulate transcription as well as gene silencing when cells are treated with CHX give the cells an advantage during Erastin stress is an interesting question.

**Figure 5:**
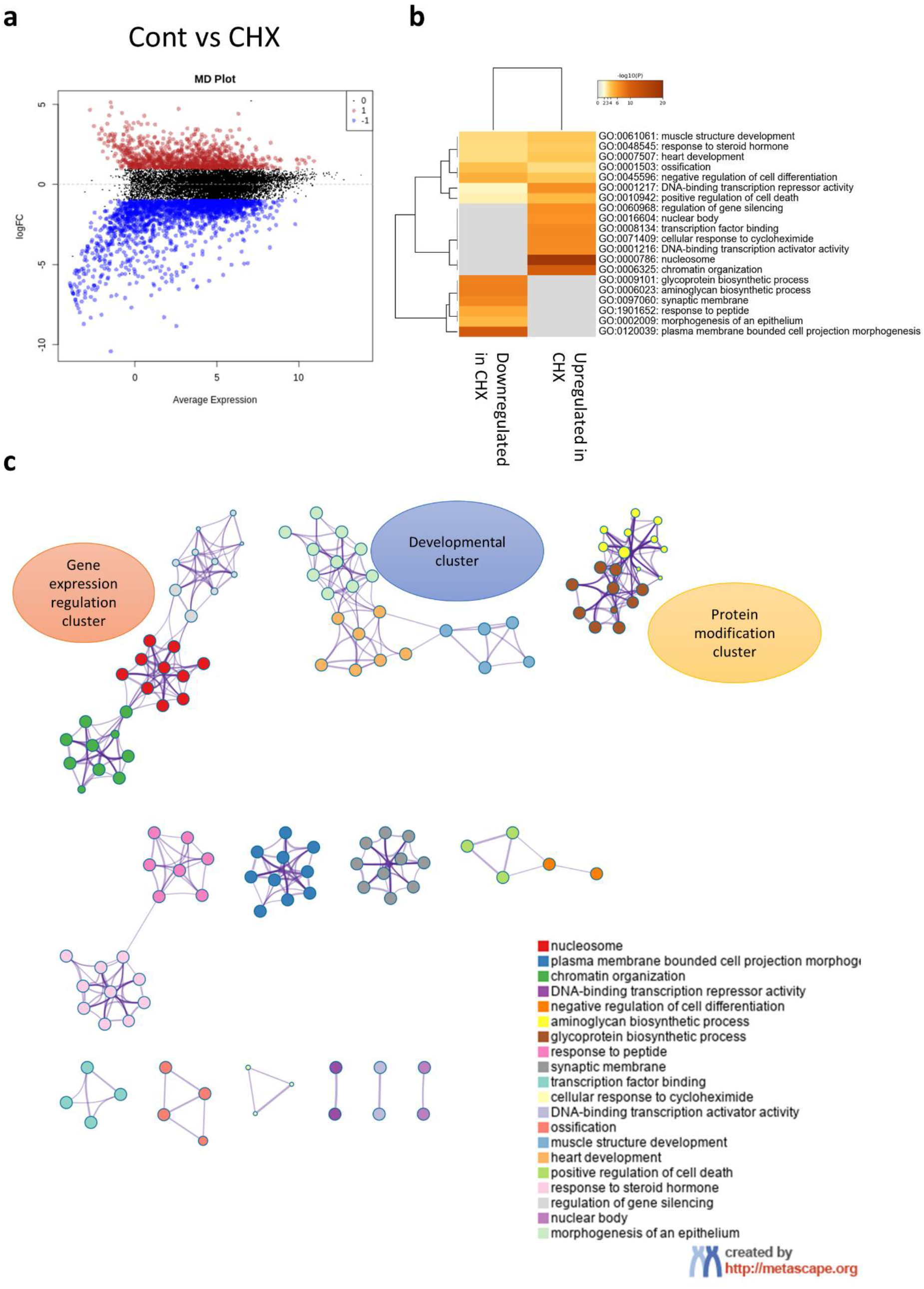
Cycloheximide alters cellular transcriptome. RNA-seq analysis of the impact of CHX on cellular transcriptome. a: MD plot revealing hundreds of DEGs following CHX treatment (4 hours treatment). b: GO analysis of differentially expressed genes after CHX treatment. c: Cluster analysis of enriched pathways reveals several important clusters related to gene expression regulation (upregulated in CHX), Developmental cluster (Mostly downregulated), and protein modifications (Downregulated). See supplementary figure 8 for more information.

### mRNA stability as a factor in ferroptosis response

ActD blocks the transcription of RNA from DNA and is frequently used to analyze mRNA stability via this property^34^. Following transcription blockade, mRNA half-life becomes a function of how effectively this mRNA is translated^35^, which is dependent on an array of epigenetic and epitranscriptional factors. In order to analyze if a different pattern of mRNA stability, and in return potential differences in translation, exist following ferroptosis, we compared each of the control, Erastin and RSL3 to their ActD treated counterpart. We hypothesized that if mRNA stability is equal between transcripts, thus we shall not observe major differences in this analysis (e.g., RLS3 vs RSL3 + ActD). If DEGs are observed, that indicates a change in the mRNA levels in relation to the global mRNA pool and can be considered as differential stabilization or destabilization of mRNAs (depending on whether they were up or downregulated) and warrants further investigation [Supplementary figure 9a]. Most genes were downregulated after ActD treatment in this analysis, which, based on our starting assumptions, indicates they are more destabilized in relation to the total mRNA pool, and only few hundred genes were upregulated, which were deemed to be stabilized [Data not shown]. To further identify the differentially stabilized or destabilized genes, we compared each dataset with the gene sets from Control vs ActD only groups. This yielded the genes that were differentially stabilized or destabilized after Erastin or RSL3 treatment (as compared to no ferroptosis stress control) [Supplementary figure 9b and c]. It was observable, however, that in the uniquely stabilized genes, one of the top terms in both Erastin and RSL3 was the polysomal ribosome. Moreover, only few terms were enriched in the Erastin treatment group as compared with the RSL3 group [Supplementary figure 9b]. In the destabilized genes, several terms were enriched in both groups, including cellular response to glucose starvation, autophagy, and response to decreased oxygen levels [Supplementary figure 9c]. However, most of the terms did not overlap. Intriguingly, glucose starvation was shown to block ferroptosis via AMPK-dependent mechanism^36^. The destabilization of genes related to cellular response to glucose starvation might thus be a mechanism by which cells respond to ferroptosis via regulating their energy expenditure. Nonetheless, this remains to be tested. Another important observation was the enrichment of the GO term “negative regulation of translation initiation” in the Erastin dataset only [Supplementary figure 9c]. Destabilization of mRNA should reflect a reduction in translation efficiency or reduced availability for translation due to their shorter half-life. One might consider the destabilization of genes that would negatively impact translation initiation to be a mechanism by which cells can regulate their mRNA translation in response to stress and, in theory, maintain an efficient translation machinery to regulate the stress response.

CHX was previously shown to selectively impact the stability of certain mRNAs^37,38^. Hypothesizing a transcriptome wide effect on mRNA stability following CHX, we repeated the same analysis as with ActD and analyzed uniquely stabilized and destabilized genes following CHX treatment in Erastin and RSL3 groups [Supplementary figure 10a and b]. In the stabilized gene dataset, we could not identify significant GO term that would correlate with the observed mechanism. In the destabilized genes, the same was true apart from 2 pathways. The “regulation of transcription from RNA polymerase II in response to stress”, enriched in RSL3, can indicate an interference with the normal RSL3 induced transcriptional stress response observed [Figure 3b], thus accentuating the cell death. The other interesting term was the tRNA metabolic process, enriched in Erastin. In theory, this can impact translation via affecting tRNA dynamics and availability. Whether this pathway contributes to the generation of GSH observed with CHX and Erastin co-treatment in a manner similar to the tRNA changes observed previously^18^ remains to be evaluated.

Collectively, the stability analysis provides clues to the influence of translation and transcription on ferroptosis response and the subtle nuances between cellular response to Erastin or RSL3. However, it appears that the global process of transcription or translation repression, rather than what genes are affected by such inhibition, is what guide the cellular fate in Erastin or RSL3.

### ALKBH1 mediated translation repression in glioma and ferroptosis sensitivity

Based on our results, we hypothesized that a repressed global protein translation cellular landscape would be beneficial against Erastin but deleterious when cells are treated with RSL3. Translation repression is known to occur in several cancers via tRNA modifications driven process^39^. Ferroptosis is important in cancer biology, with many aspects of tumor behavior and microenvironment heavily dependent on the tumor susceptibility or resistance to ferroptosis^40,41^. Moreover, several anti-cancer therapies for ferroptosis are being considered^42^. Thus, understanding the nuances of tumor response to ferroptosis inducing agents is paramount in the quest to treat various cancers. To that end, we chose Alkbh1 as a target gene for our evaluation. Alkbh1 is associated with worse outcome in Glioblastoma (GB) patients^43^. Alkbh1 is a tRNA demethylase that acts on 1-methyladenosine (m^1^A) leading to tRNA hypomodifications, reduced tRNA stability, and translation repression^10,44^.

To test for our hypothesis, we generated cells overexpressing Alkbh1 using lentiviral transfection of rat 9L gliosarcoma cell line, which simulates high grade glioma [Figure 6a]. Cells overexpressing Alkbh1 had slower growth rates and protein translation rates, which are consistent with its previously observed tRNA demethylation activity^44^ [Figure 6b and c]. Alkbh1 overexpressing cells were more resistant to complex III inhibition via Antimycin A (contrary to previous observation^10, 45^) [Figure 6d] and had paradoxical faster migration rates than control cells under normal conditions and in the presence of mitochondrial oxidative stress inducing agents that are known to induce mitochondrial dysfunction induced senescence (MIDAS)^46^; Antimycin A and Rotenone [Figure 6e, supplementary figure 11]. Alkbh1 mediated resistance to MIDAS was more evident with Rotenone induced mitochondrial dysfunction, in-line with the role of Alkbh1 in regulating respiratory complex I activity^45^.

**Figure 6:**
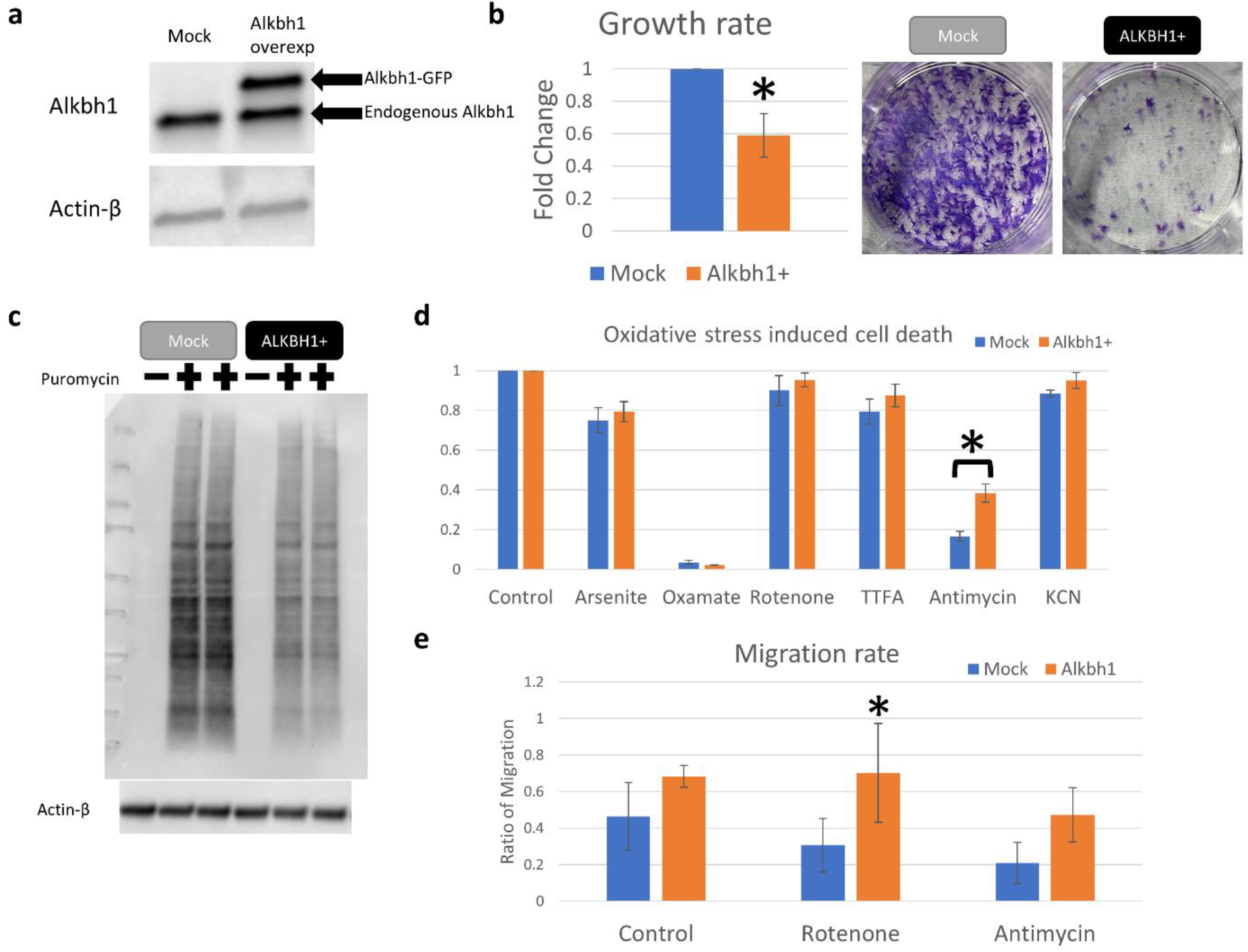
Alkbh1 overexpression in glioma cells alters growth and translation. a: Confirmation of Alkbh1 expression in 9L gliosarcoma cells using western blotting. Alkbh1 fused with GFP was robustly detected. b: Alkbh1 overexpressing cells (Alkbh1+) had slower growth rate as evident by WST-8 assay (Graph) and Cresol violet staining (Photos). c: Western blot analysis of puromycin incorporation assay reveals reduced global protein translation in Alkbh1+ cells. d: Glioma cells were exposed to toxic doses of Arsenite (endoplasmic reticulum stress, non-specific), sodium Oxamate (LDH inhibitor), Rotenone (Mitochondrial respiratory Complex I inhibitor), TTFA (Complex II), Antimycin (Complex III), and KCN (potassium cyanide; Complex IV). Alkbh1+ cells were more resistant to Complex III inhibition. e: Analysis of cell migration using Scratch test shows that Alkbh1+ cells migrate faster than Mock cells, this is especially evident when Rotenone was added.

Next, we analyzed genes that can explain the observed behavior of Alkbh1 overexpressing cells or the cause of worsened outcome in glioblastoma with high Alkbh1 expression. qPCR analysis revealed upregulation of Oct4, the stemness marker which is associated with enhances proliferation and colony formation in glioma cells^47^ [Supplementary figure 12a]. Sox2 and Sox4 expression did not change with Alkbh1 overexpression [Supplementary figure 12a]. Alkbh1 is known to impact mitochondrial activity and to localize into the mitochondria^10,45,48^. To evaluate its impact on glioma mitochondrial function we evaluated several mitochondrial Sirtuin genes expression levels [Supplementary figure 12b]. Only Sirt5 was upregulated after Alkbh1 overexpression. Sirt5 is associated with more resistance to MIDAS^46^ and can support tumorigenesis in breast cancer^49^. Examination of the cytosolic and mitochondrial chaperones revealed strong upregulation of Hsp70 and Lonp1 [Figure 12c], which signifies activation of the unfolded protein responses and better handling of misfolded proteins^50,51^. Hsp70 plays important role on glioma pathogenesis via various mechanisms including apoptosis evasion, angiogenesis, and supporting mutated proteins and genome instability^52^. Lonp1 also plays important role in glioma and is associated with worse outcome^53^. MMP2 expression was also upregulated signifying enhanced tumor invasion^54^ [Supplementary figure 12d]. Finally, we evaluated genes that are associated with glioma migration. Stx1a, Doublecortin (Dcx), and Ttyh1 were extremely upregulated after Alkbh1 expression. Interestingly, these genes are associated not only with glioma migration and invasion, but also with synaptic development and intercellular communication and neurotransmitter signaling^55^. This can represent an interesting mechanism by which Alkbh1 promotes glioma progression via activating the neurogenic pathways and synaptic development^56^. Overall, this data indicates that Alkbh1 supports glioma progression and invasion via multiple mechanisms including better proteostasis in the cells via protein translation repression and activation of the heat shock and chaperone proteins, promoting invasiveness, activation of mitochondrial Sirtuin network, and promoting synaptic formation.

mRNA sequencing revealed hundreds of genes to be differentially expressed in Alkbh1 cells vs Mock transfected cells [Figure 7a]. GO analysis revealed a significance downregulation of pathways related to DNA replication and cell division and upregulation of pathways related to cellular immune responses [Figure 7b and supplementary figure 13]. Analysis of transcription factors motifs (TFs) that might be responsible for the gene expression pattern revealed E2F1 and HIF-1α motifs are the top enriched in the downregulated genes while RELA/P65 and NF-κβ targets are most enriched in upregulated genes [Figure 7c]. E2F1 and HIF1A are associated with reduced migration and tumorigenesis in glioma^57,58^. Downregulating gene targets of both TFs signifies enhanced invasion and migration of glioma cells, in concordance with our observations. Activation of NF-κβ/RELA/p65 pathway is on the other hands associated with enhanced migration, stemness, proinflammatory response in tumor, and resistance to therapy^59–61^. This is also in-line with our observations above. Finally, protein-protein interaction (PPI) network analysis was performed. PPI network analysis confirmed the activation of inflammatory and immune pathways in Alkbh1 overexpressing cells as well as enhanced glioma migration and suppression of DNA replication [Supplementary figure 14, supplementary table 3]. Alternative splicing analysis revealed enrichment and depletion of hundreds of AS events in Alkbh1 overexpressing cells. The most significant AS program was skipped exon (SE), while other types showed relatively equal number of LSV events [Supplementary table 4]. GO analysis revealed several pathways related to protein translation and quality control as well as histone and chromatic related activities to be highly enriched in AS dataset [Figure 7d]. Overall, Alkbh1 drives a transcriptional and epigenetic program that promotes glioma migration, invasion, immune modulation, and stemness while suppressing DNA replication and cell division at the same time.

**Figure 7:**
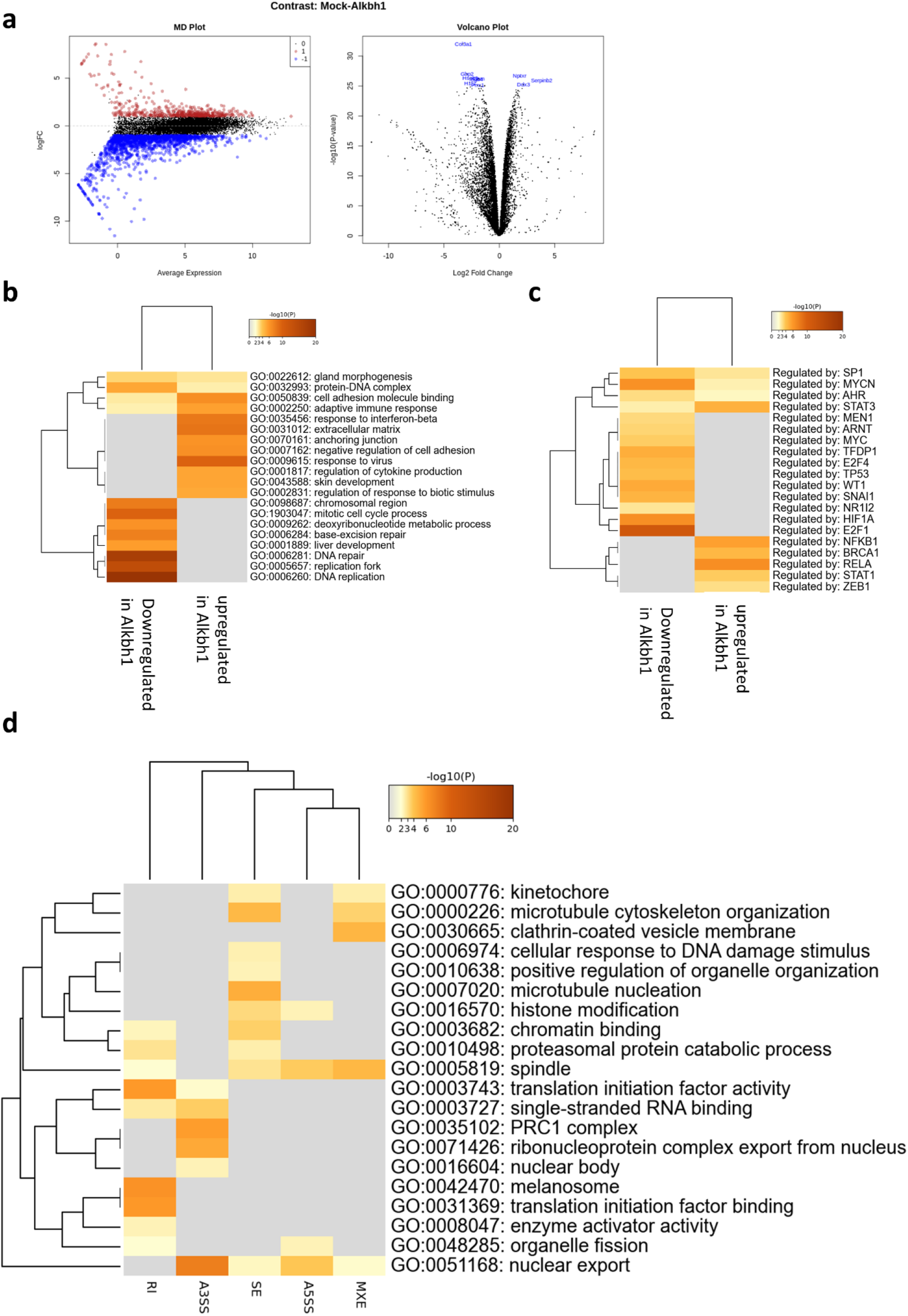
RNA-seq analysis of Alkbh1 impact on cellular transcriptome. a: MD plot (left) and volcano plot (right) showing hundreds of DEGs between Mock and Alkbh1+ cells. b: GO analysis reveals downregulation of pathways related to DNA replication and cell growth and upregulation of immune-related pathways. c: Transcription factors (TF) analysis reveals that the downregulated genes are mostly regulated via E2F1 and HIF-1α, while the upregulated genes are regulated via NF-κβ/RELA system.

Finally, to test our hypothesis, cells were exposed to Erastin or RSL3 and cell death observed. Alkbh1 overexpressing cells were more resistant to Erastin induced ferroptosis but were more sensitive to RSL3 induced ferroptosis [Figure 8a-c]. This observation supports our hypothesis that cells with global translational repression would be more sensitive to RSL3 but more resistant to Erastin. We further queried important effectors in cellular sensitivity to ferroptosis via qPCR. Alkbh1 led to enhanced expression of Slc7a11 and Slc3a2, both members of the system xCT [Figure 8d]. This can further explain the resistance of Alkbh1 overexpressing cells to Erastin induced ferroptosis^62^. Ferroptosis suppressing protein 1 (FSP1, formerly known as Aifm2^7^) was also upregulated in Alkbh1 overexpressing cells. FSP1 is known to protect against RSL3 induced ferroptosis^7^. However, this appeared to be countered by the effect of the global translational landscape of Alkbh1 overexpressing cells, with a net outcome of more sensitivity to RSL3 induced ferroptotic cell death. Collectively, Alkbh1 protects glioma cells against Erastin induced ferroptosis via protein translation repression as well as via activating members of the system xCT. The sensitivity of Alkbh1 overexpressing cells to RSL3 is, at least to a large part, derived by the translation repressive effect of Alkbh1.

**Figure 8:**
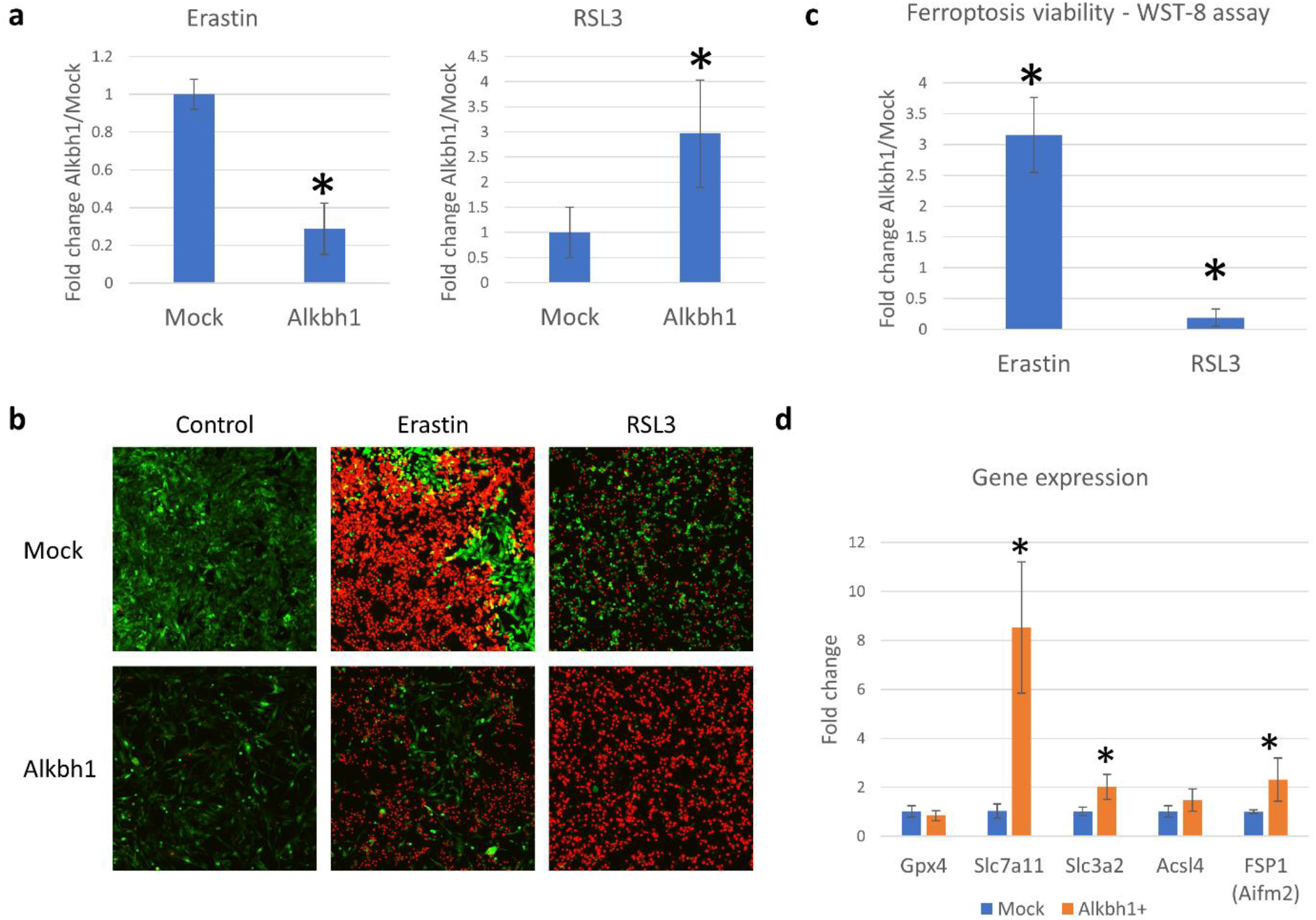
Alkbh1 protects glioma cells from Erastin induced ferroptosis but make them vulnerable to RSL3 induced ferroptosis. Cells were stress with Erastin (5μM, 24 hours) or RSL3 (1μM, 6 hours). a: Analysis of PI staining for dead cell counting shows much lower dead cell count in Alkbh1+ cells after Erastin treatment with reversal of that effect after RSL3 treatment. Asterisk = statistically significant (*p* < 0.0005) versus Mock. b: Example of PI staining. c: WST-8 assay showing the difference in cell viability between Alkbh1+ cells and Mock cells after Erastin or RSL3 treatment. Asterisk = statistically significant (*p* < 0.0005). d: Gene expression analysis of important genes known to impact ferroptosis sensitivity. Asterisk = statistically significant (*p* < 0.05, Fold change > 1.5) versus Mock.

## Discussion

The results presented in this work are multifold; first, we identified a strong AS program that is activated earlier and to a greater extent than transcriptional changes during cell response to ferroptosis. Importantly, the landscape of this AS program is different between Erastin and RSL3 induced ferroptosis. Further supporting the notion that while both drugs have an end phenotype of ferroptotic cell death, the road leading to this end point is vastly different epigenetically as well as metabolically^14^. AS is regulated by several mechanisms, including RNA binding proteins (RBPs)^63^ and RNA modifications^64^. Recently, several RBPs were shown to regulate ferroptosis^65,66^. For example, ELAVL1 knockdown, which is known to impact AS of different mRNAs^67^, was shown to protect against Erastin^66^. However, a global understanding of the functions of different RBPs and the role of Alternative splicing remains largely unexplored in ferroptosis.

The second aspect of our findings is identifying the role of mRNA translational landscape in dictating cellular fate in ferroptosis, and how it differentially impacts ferroptosis induced by system xCT or GPX4 inhibition. Translation repression is an important mechanism in cellular stress response, and it involves a myriad of mechanisms that coordinate changes aiming to protect cells^68^. One of the important mechanisms involved in regulating mRNA translation is m6A mRNA modifications^69^. mRNA modifications also have important role in AS^64^. Stress induced translation repression is indeed a dynamic mechanism. Activation of different stress-activated kinases that block translation initiation^68^, stress granule formation and sequestration of mRNAs from translation sites^70^, tRNA transcript expression changes that alter the availability of amino acids for translation^18^, tRNA cleavage and tRNA derived small RNAs^71^, and other non-coding RNAs mediated translation repression^72^ are all mechanisms that contribute to stress induced translation repression. Most of these mechanisms are not explored nor focused on in the context of ferroptosis. Importantly, many of these mechanisms play important role in many diseases that has links to ferroptosis. For example, stress granules play important role in Alzheimer’s disease and other neurodegenerative disease^73^, where ferroptosis also plays important role. tRNA modifications also play important roles in many cancers^39,74^. Given the importance of ferroptosis in cancer biology, it will be insightful to establish any potential links between these two mechanisms as highlighted in our analysis of Alkbh1 links to ferroptosis.

Recently, a link between the integrated stress response (ISR) and Erastin induced ferroptosis was established^12^. Knockdown of MESH1 led to activated of ISR, endoplasmic reticulum (ER) stress, phosphorylation of eIF2α and protected against Erastin induced ferroptosis^12^. eIF2α phosphorylation suppresses global mRNA translation^75^. Such global translation suppression is associated by selective translation of stress responsive genes such as GSH^18^ via mRNA stabilization and selective tRNA enrichment^18^. In our work we observed a similar mechanism when CHX was used alone or with ferroptosis inducers. CHX led to upregulation of GSH, which means that CHX did not block all translation, but acted in a way similar to ISR, at least in terms of few hours treatment. This observation is important to understand the impact of mRNA translation repression in ferroptosis and how it modulates the response to different ferroptosis inducing drugs classes. It also has a very important technical implication. CHX is used to block translation during Ribosome profiling experiments to probe differentially translated mRNAs and codon enrichment^76^. CHX was shown to stimulate the transcription of genes involved in ribosomal biogenesis in yeast in a study that cautioned on interpreting the results of ribosome profiling experiments^31^. Our results reveal that stress responsive mRNAs might escape CHX mediated translation repression, which might also have an impact on the interpretation of ribosome profiling experiments. We believe our results are basis to justify further examination of this effect and how it might distort the interpretation of ribosome profiling results.

The third aspect is providing new information on the link between CHX and the activation of the reverse transsulfuration pathway in the protection against Erastin induced ferroptosis. Transsulfuration pathway is a well-known process for the regeneration of GSH^77^. Thus, in translationally repressed cells, a shift to GSH generation from Homocysteine can accommodate for the loss of Cystine, which is a consequence of Erastin induced inhibition of system xCT^8,23,24^. Interestingly, this mechanism appears to not be functioning in transcription repression mediated blockade of Erastin induced ferroptosis, nor it has any bearing on RSL3 induced ferroptosis.

The fourth aspect of our results is identifying that translation repression mediated via certain tRNA modifications (in our case m^1^A demethylation via Alkbh1) can be used to identify synthetic lethality of cancer cells to different ferroptosis classes. This opens the doors to the utilization of tRNA epitranscriptome as biomarkers for personalizing ferroptosis therapy. Alkbh1 overexpression worsen the survival in glioblastoma patients^43^. We argue that identification of the molecular signature of Alkbh1 activity, such as reduced tRNA methylation^10,44^, can be used as biomarkers to adjust the therapy in these patients. We also believe that this mechanism is not limited to Alkbh1 nor to m^1^A. Many tRNA modifications can impact mRNA translation, either by impacting codon-anticodon pairing (in case of modifications at the anticodon), or by altering tRNA conformation and stability (such as with methylation modifications not in the anticodon)^39,71,74^. Systematic analysis of the links between tRNA epitranscriptome, mRNA translation, and ferroptosis would ultimately lead to great insights into these interesting mechanisms.

Finally, our results could not accurately pinpoint the exact mechanism by which ActD-mediated transcription repression blocked Erastin induced ferroptosis. Theoretically, changes at the epigenome and epitranscriptome level would alter the translation rates of mRNAs leading to changes in mRNAs half-lives and stability^38,74^. While our data could give crude overview of the global changes in mRNA stability in relation to the global pool of RNA, it is not precise measurements. Future work using more precise ways to measure mRNA decay rates^78^ during ferroptosis, as well as transcription rates, would indeed provide the insight needed to understand the link between transcription repression and ferroptosis as well as the various epigenetic mechanisms governing mRNA stability during cellular stress response to lipid peroxidation.

In summary, our results reveal an important, yet not fully explored, role of the epitranscriptome and epigenome in dictating cellular response and sensitivity to ferroptosis inducing agents. Our work also reveals important differences between various classes of ferroptosis inducing agents and their links to the transcriptional, translational, and epigenetic landscape in the cell that can be used to further understand the process of ferroptosis or identify novel mechanisms for disease therapies. We hope our work can serve as basis for future exploration of these mechanisms and provide foundations for studying the epigenome and epitranscriptome in ferroptosis.

## Methods

### Cell culture

B35 neuroblastoma cells were purchased from ATCC (Cat# CRL-2754, RRID: CVCL_1951). 9L gliosarcoma cells were a gift from Dr Ichiyo Shibahara (ATCC). Both cell lines were cultured in high glucose DMEM (Nacalai Tesque, Cat# 08457–55) containing 10% fetal bovine serum (FBS; Hyclone, GE-life sciences, USA. Cat# SH30910.03 for 9L cells or Corning, Cat# 35-079-CV for B35 cells). All B35 experiments were conducted on Poly L-lysine (PLL) coated plates and dishes. 9L experiments were conducted on non-coated plates and dishes.

### Reagents and drugs

Erastin (Sigma, Cat# E7781). 1S-3R-RSL3 (RSL3, Sigma, Cat# SML-2234). Actinomycin D (ActD, Sigma, Cat# A9415), Cycloheximide (CHX, Sigma, Cat# C7698). Bodipy 581/591 C11 (Invitrogen, Cat# D3861). Cell count reagent SF (For WST-8 assay, Nacalai tesque, Cat# 07553-44). Calcein green (Invitrogen, Cat# C34852). Puromycin (Sigma, Cat# P8833). Ferrostatin-1 (Ferr-1, Sigma, Cat# SML0583). Sodium meta-Arsenite (Arsenite, Sigma, Cat# S7400). Rotenone (Sigma, Cat# R8875). Antimycin A (Sigma, Cat# A8674). Potassium Cyanide (KCN, Sigma, 60178). Sodium oxamate (Oxamate, Sigma, Cat# O2751). 2-Thenoyltrifluoroacetone (TTFA, Sigma, Cat# T27006).

### Western blotting

Western blotting was performed as previously described^10^. In brief, Cells were homogenized in protein extraction buffer containing Triton-x, centrifuged, and the supernatant containing proteins separated. Protein concentrations were evaluated using Brachidonic acid assay kit (Thermo Fisher Scientific, Cat# 23227). Equal loads of proteins were separated on Mini-PROTEAN TGX gel (Bio-rad) and transferred to polyvinylidene difluoride membrane (Biorad) using semi-dry transfer. Membranes were blocked in 5% skim milk solution in PBS-T, incubated with primary antibody of choice overnight at 4°C, incubated with secondary antibody (IgG detector solution) and the signal detected using chemiluminescence Western blotting detection reagents (Pierce ECL Western Blotting Substrate, Thermo Fisher Scientific Inc., Rockford, IL, the USA, Cat 32106) on ChemiDoc machine (Bio-Rad). Membranes were stripped and re-probed using anti-beta actin antibody as a loading control.

Antibodies used were anti-Alkbh1 antibody (1:500 dilution, Abcam, Cat# ab195376). anti-beta actin antibody (1:5000 dilution, Cell signalling technologies, Cat# 4970 S). anti-puromycin antibody (1:2000 dilution, Millipore, Cat# MABE343). Secondary antibody used was IgG detector solution, HRP-linked (1:2000, Takara, Japan, Cat# T7122A-1).

### Puromycin assay

Cells were incubated in full growth medium containing 10μg/ml puromycin for 30 minutes, the collected as explained above and analyzed using western blotting using anti-puromycin antibody as demonstrated above.

### Generation of stable Alkbh1 overexpressing glioma cell line

Alkbh1 Rat Tagged ORF Clone Lentiviral Particles and Mock control were purchased from Origene (Cat# RR214755L2V). Cells were transfected with lentiviral particles in the presence of polybrene (Sigma, Cat# TR-1003-G) as previously described and colonies selected via limiting dilution and single colony expansion^10^. Colonies were monitored and selected on the basis of their GFP expression. Following selection and expansion, western blotting was used to evaluate expression of exogenous Alkbh1, and to evaluate the stability of transfection over multiple passages. Multiple clones were successfully established, and one was selected at random for the purpose of this work.

### Evaluation of cell death and ferroptosis

Cell viability after exposure to ferroptosis inducing agents was evaluated via different methods such as WST-8 assay to evaluate cell viability, Live cell imaging by Calcein green for living cells, and Propidium iodide (PI) staining for dead cells. Number of dead cells was counted using ImageJ software and presented a ratio between Alkbh1 overexpressing cells and Mock cells. Bodipy C11 was used to evaluate lipid peroxidation by observing the shifts in fluorescence emission as described previously^79^.

### Analysis of cell growth and migration

Analysis of cell growth in Alkbh1 overexpressing 9L cells or Mock cells was performed by platting the cells at very low confluence (1000 cells/ well in 96 or 6 well plates). Cells were grown for 1 week and analyzed using WST-8 assay (96 wells), or grown for 2 weeks, then fixed, and stained using Cresol violet (6 wells).

Analysis of glioma migration was performed using scratch assay. Cells were plated in 6 well plates 24 hours before the test in order to be >90% confluent by the time of the test. A scratch was performed using a 200μl pipette tip. The width of the gap was measured immediately after scratch (0 hour), and cells incubated for 24 hours in fresh growth medium, then the gap width was measured again. The ratio of migration was calculated using this equation:

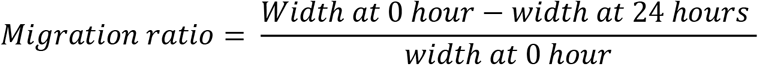

### UHPLC-MS/MS analysis

UHPLC-MS/MS was performed as detailed previously^80^. Samples were prepared by plating equal number of B35 cells in 6 well dishes (1 × 10^6^/well) and exposing the cells to Erastin (1μM) or RSL3 (0.1μM) with or without ActD (5μg/ml) or CHX (100μg/ml) for 8 hours. Cells were washed in ice cold PBS and suspended in 100μl methanol containing internal standards, homogenized by sonication, deproteinized and analyzed by ultra HPLC triple quadrupole MS (UHPLC-MS/MS). The analysis was performed on an Acquity™ Ultra Performance LC I-class system (Waters Corp., Milford, UK) interfaced to a Waters Xevo TQ-S MS/MS system equipped with electrospray ionization. Details of the analysis were published previously^80^. The data were collected using the MassLynx version 4.1 software (Waters Corp.) and the ratio of the peak area of analyte to the internal standard was analyzed by Traverse MS (Reifycs Inc., Tokyo, Japan)^80^. The analysis was performed using 6 samples per group and data presented as Area/ratio.

### RNA-sequencing

Cells were lysed in Qiazol (Qiagen, Cat# 79306) following exposure to drugs as indicated (B35 ferroptosis experiment) or without drug treatment (9L Alkbh1 overexpression experiment). RNA was extracted using Qiagen’s miRNeasy mini kit (Cat# 217084) with DNase digestion step. RNA concentration and purity was determined using nanodrop 1 (ThermoFisher). RNA integrity was analyzed using Agilent Bioanalyzer 2100 and Agilent’s RNA 6000 nano kit (Cat# 5067-1511). RNA integrity number (RIN) was above 9 in all samples used. mRNA sequencing was performed using NEBNext Ultra II Directional RNA Library Prep kit (Cat# E7765S) as per manufacturer’s instructions. Quality control of libraries was performed using Agilent DNA 1000 kit (Cat# 5067-1504). NEBNext Library Quant Kit (Cat# E7630L) was used for quantifying libraries. Libraries were pooled and sequenced on Hiseq-X ten (150bp, paired end) by Macrogen Japan. All sequencing experiments were performed with 3 replicates per group.

### RNA-sequencing data analysis

Quality control for Raw fastq files was performed using FastQC. Reads were trimmed using Trimmomatic^81^ and adaptor sequences and low-quality reads removed. Reads were aligned to Rattus norvegicus reference genome (mRatBN7.2/rn7 assembly from UCSC) using Hisat2^82^ with splice aware options using GTF file for reporting. Read counting and transcript assembly were performed using Stringtie^83^, and differential gene expression was performed using Limma-voom package^84^. Fold change > 1.5 (in the B35 ferroptosis experiment) or > 2 (in the 9L Alkbh1 overexpression experiment) plus FDR < 0.05 were criteria for statistical significance of differentially expression genes (DEGs).

Alternative splice (AS) analysis was performed using the rMATs-turbo suite^28^. Statistical significance of alternative splicing events was considered when FDR < 0.05 and delta-Psi (ΔΨ) > 0.2.

Gene ontology and pathway analysis was performed using Metascape^85^. Only genes satisfying the selection criteria in DEGs analysis or AS analysis were selected for pathway analysis.

### Real-time PCR (qPCR)

RNA was extracted as above, and cDNA synthesized from 1μg RNA using Superscript III reverse transcriptase (ThermoFisher, Cat# 18080044). qPCR was performed using GoTaq qPCR Master Mix (Promega, Cat# A6102) on CFX96 thermal cycler (Bio-Rad). qPCR was conducted with 4 replicates per group. qPCR was analyzed using the ΔΔCT method with Actin-β as an internal control.

Primers used in this study:

Actin-β: Forward: 5’GGAGATTACTGCCCTGGCTCCTA,

Reverse: 5’GACTCATCGTACTCCTGCTTGCTG

Acsl4: Forward: 5′AATTCCACCCTGATGGATGCTTAC, Reverse: 5′TTCAGTGCGGCTTCGACTTTC

FSP1 (Aifm2): Forward: 5′CCATCCAGGCCTATGAGGACA, Reverse: 5′TTAATCTCTGCGGCCATCTCAAC

Dcx: Forward: 5′CACTGACATCACAGAAGCGATCAA, Reverse: 5′TCAGGGCCACAAGCAATGAA

Gja1: Forward: 5′AGGTCTGAGAGCCTGAACTCTCATT Reverse: 5′GGCACTCCAGTCACCCATGT

Gpx4: Forward: 5′ATGCCCACCCACTGTGGAA, Reverse: 5′GGCACACACTTGTAGGGCTAGAGA

Hsp70: Forward: 5’GCTCGAGTCCTACGCCTTCAATA, Reverse: 5’CTCAGCCAGCGTGTTAGAGTCC

Hspa9 (GRP75): Forward: 5’CCGGAGACAACAAACTTCTAGGACA, Reverse: 5’TTGGCAGAAACGTGCACAATC

Hspd1 (HSP60): Forward: 5’ACCGGAAGCCCTTGGTCATAA, Reverse: 5’TGGAGCTTTGACTGCTACAACCTG

Lonp1: Forward: 5′TGACAGATGTGGCAGAAATCAA, Reverse: 5′GGTAGCCTCGGCCAATCTTA

MMP2: Forward: 5’TCCCGAGATCTGCAAGCAAG Reverse: 5’AGAATGTGGCCACCAGCAAG

MMP9: Forward: 5’AGCCGGGAACGTATCTGGA Reverse: 5’TGGAAACTCACACGCCAGAAG

Oct4: Forward: 5′CATCTGCCGCTTCGAG, Reverse: 5′CTCAATGCTAGTCCGCTTTC

Sirt1: Forward: 5′CGGACAGTTCCAGCCATCTCTG, Reverse: 5′GGATTCCTGCAACCTGCTCCAA

Sirt3: Forward-5′AAGACATACGGGTGGAGCCT Reverse-5′GGACTCAGAGCAAAGGACCC

Sirt5: Forward-5′TTACCACTACCGGAGGGAGG Reverse-5′TGATGACCACAACCCGTCTG

Slc3a2: Forward: 5′CACTCCCAACTATAAGGGCCAGAA, Reverse: 5′CATCCCGAACTTGGAAACCATC

Slc7a11: Forward: 5′GTGGGCATGTCACTGGTGTTC, Reverse: 5′GCTCGTACCCAATTCAGCATAAGA

Sox2: Forward: 5′GTCAGCGCCCTGCAGTACAA, Reverse: 5′GCGAGTAGGACATGCTGTAGGTG

Sox4: Forward: 5′TGGACGTATTTATACTGGCCAAACA, Reverse: 5′AATGGCGTGGGTAACAACACTAGA

Stx1a: Forward: 5′GGCCGTCAAGTACCAGAGCAA, Reverse: 5′ATGATGATGCCCAGAATCACACA

Ttyh1: Forward: 5′GACTTCTGCTCCAACCCAGACAC, Reverse: 5′AGCACGCTGGGACAGAGTCA

### Statistical analysis

Statistical analysis was performed using SPSS v20 (IBM). All data are represented as mean ± standard deviation (SD). ANOVA with Turkey post Hoc was used to compare multiple groups and independent sample Student’s t-test was used to compare between 2 groups. *p* < 0.05 was considered to be statistically significance in addition to fold change (FC) > 1.5 when appropriate. All experiments were conducted in triplicates or more.

## Data availability

Raw sequencing data were deposited at the sequence read archive (SRA) project numbers PRJNA778943 and PRJNA779470. All other data are presented in the manuscript and supplementary figures.

## Author contribution

**S.R:** Study conception and design. Administration. Funding. Conducted experiments. Analyzed the data. Wrote the manuscript. **D.S:** Conducted experiments. Funding. reviewed the final version of the manuscript. **Y.Z:** Conducted experiments, reviewed the final version of the manuscript. **L.Z:** Conducted experiments, reviewed the final version of the manuscript. **T.T:** Critically reviewed the manuscript. **K.N:** Administration. Critically reviewed the manuscript.

