## Supplementary figures for "Epigenetic regulation of translation repression in ferroptosis, and a role of Alternative splicing and tRNA methylation"

**Supplementary figure 1:** Mass spectrometry analysis (Continued from figure 2).

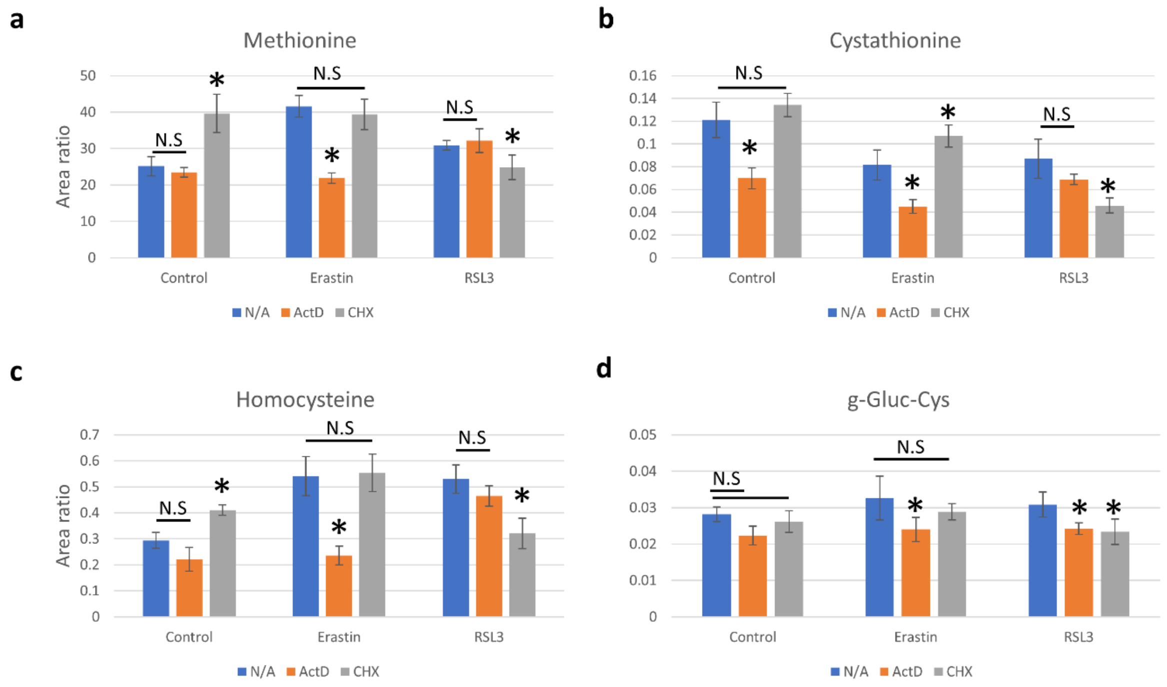

**Supplementary figure 2:** Volcano plots showing enrichment of various AS types in Erastin or RSL3 treatment groups. a: A5SS: Alternative 5' splice sites. b: A3SS: Alternative 3' splice sites. c: MXE: mutually exclusive exons. (Continued from figure 3).

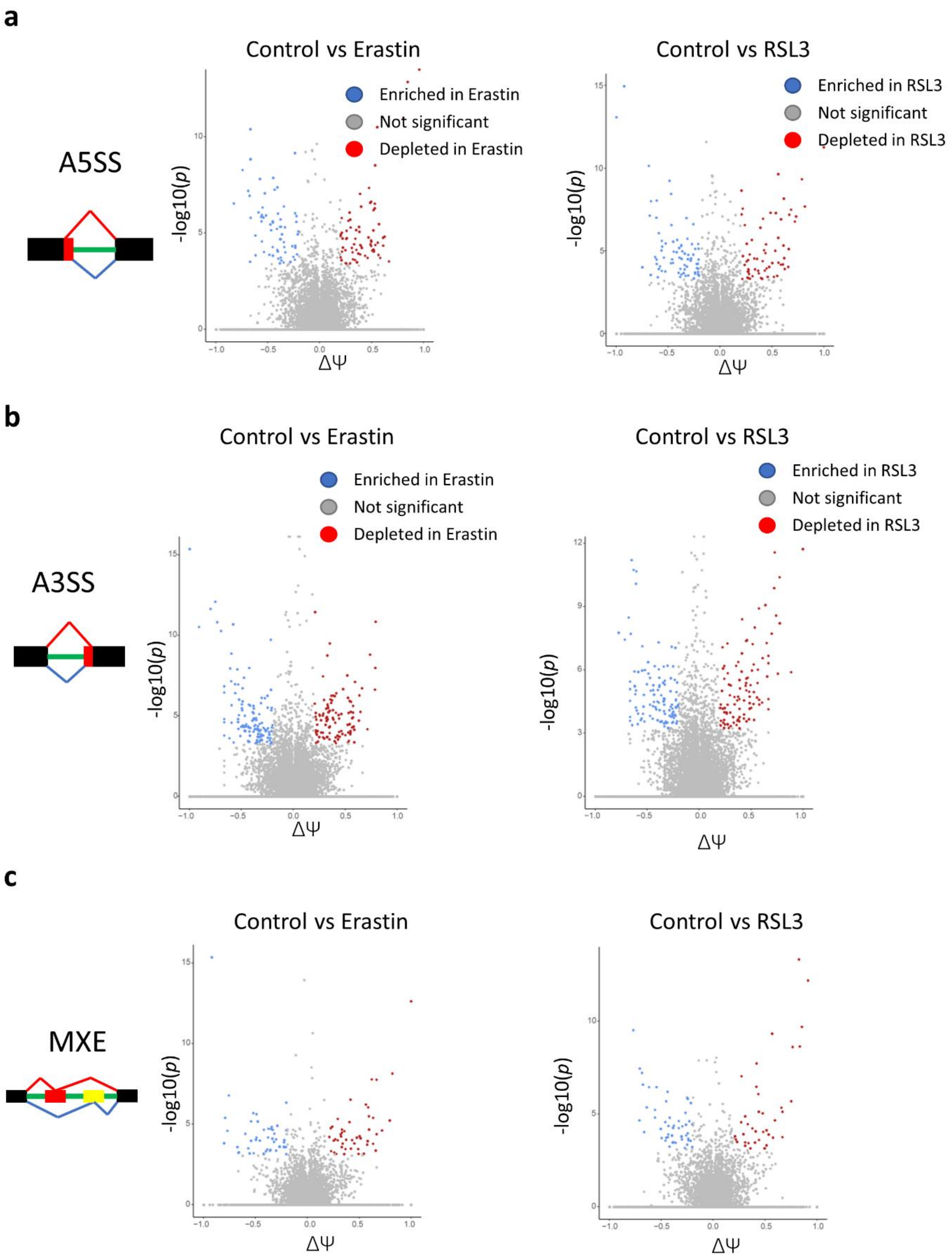
a
Control vs Erastin
Control vs RSL3
A5SS
Enriched in Erastin
Not significant
Depleted in Erastin
Enriched in RSL3
Not significant
Depleted in RSL3
-log10(p)
ΔΨ
b
Control vs Erastin
Control vs RSL3
A3SS
Enriched in Erastin
Not significant
Depleted in Erastin
Enriched in RSL3
Not significant
Depleted in RSL3
-log10(p)
ΔΨ
c
Control vs Erastin
Control vs RSL3
MXE
-log10(p)
ΔΨ

**Supplementary figure 3:** Extended GO analysis of the Erastin AS dataset showing predominance of RI events in regulating multiple pathways. (Supplemental to figure 4c)

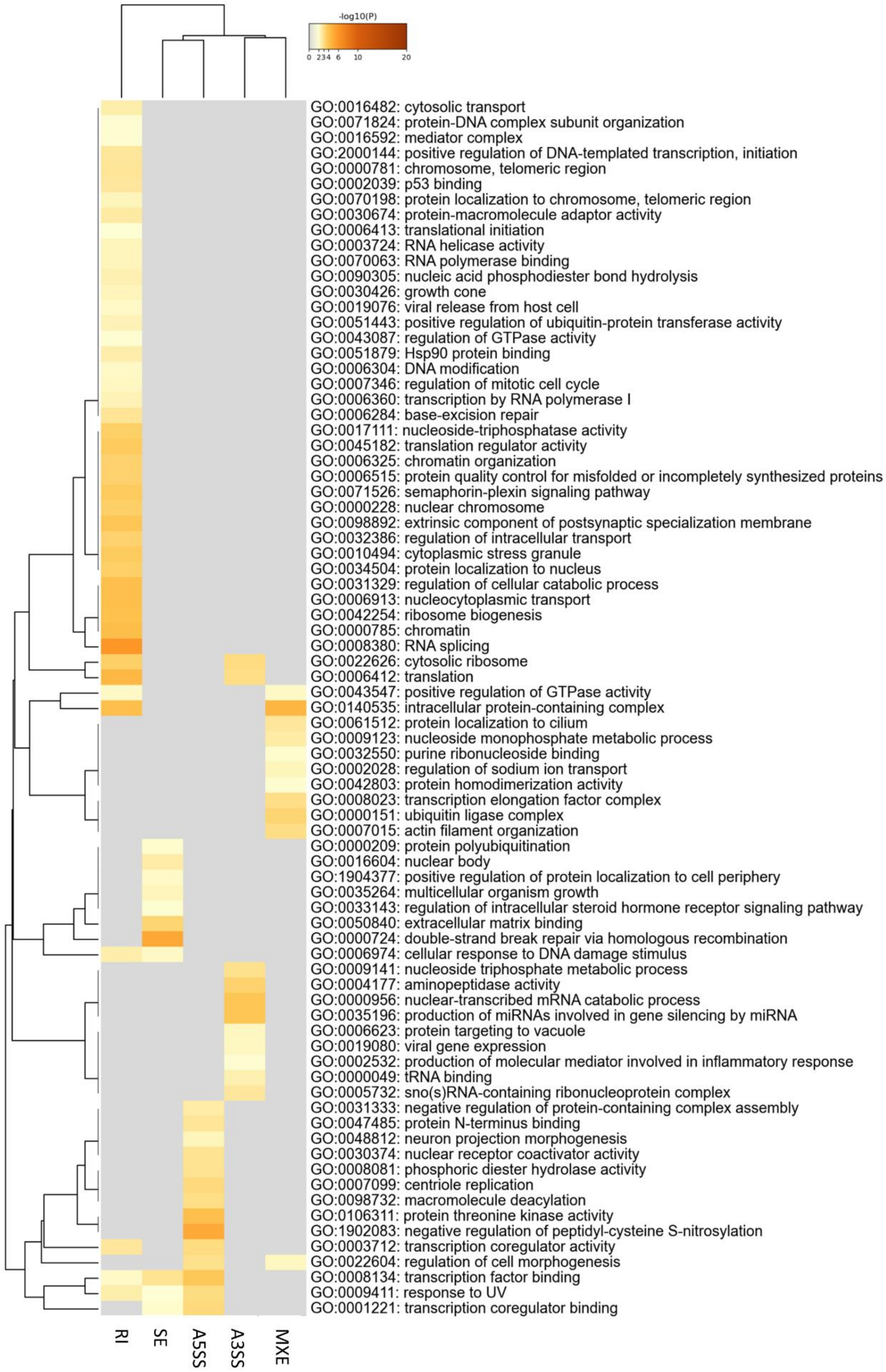
-log10(P)
0 2 3 4 6 10 20
GO:0016482: cytosolic transport
GO:0071824: protein-DNA complex subunit organization
GO:0016592: mediator complex
GO:2000144: positive regulation of DNA-templated transcription, initiation
GO:0000781: chromosome, telomeric region
GO:0002039: p53 binding
GO:0070198: protein localization to chromosome, telomeric region
GO:0030674: protein-macromolecule adaptor activity
GO:0006413: translational initiation
GO:0003724: RNA helicase activity
GO:0070063: RNA polymerase binding
GO:0090305: nucleic acid phosphodiester bond hydrolysis
GO:0030426: growth cone
GO:0019076: viral release from host cell
GO:0051443: positive regulation of ubiquitin-protein transferase activity
GO:0043087: regulation of GTPase activity
GO:0051879: Hsp90 protein binding
GO:0006304: DNA modification
GO:0007346: regulation of mitotic cell cycle
GO:0006360: transcription by RNA polymerase I
GO:0006284: base-excision repair
GO:0017111: nucleoside-triphosphatase activity
GO:0045182: translation regulator activity
GO:0006325: chromatin organization
GO:0006515: protein quality control for misfolded or incompletely synthesized proteins
GO:0071526: semaphorin-plexin signaling pathway
GO:0000228: nuclear chromosome
GO:0098892: extrinsic component of postsynaptic specialization membrane
GO:0032386: regulation of intracellular transport
GO:0010494: cytoplasmic stress granule
GO:0034504: protein localization to nucleus
GO:0031329: regulation of cellular catabolic process
GO:0006913: nucleocytoplasmic transport
GO:0042254: ribosome biogenesis
GO:0000785: chromatin
GO:0008380: RNA splicing
GO:0022626: cytosolic ribosome
GO:0006412: translation
GO:0043547: positive regulation of GTPase activity
GO:0140535: intracellular protein-containing complex
GO:0061512: protein localization to cilium
GO:0009123: nucleoside monophosphate metabolic process
GO:0032550: purine ribonucleoside binding
GO:0002028: regulation of sodium ion transport
GO:0042803: protein homodimerization activity
GO:0008023: transcription elongation factor complex
GO:0000151: ubiquitin ligase complex
GO:0007015: actin filament organization
GO:0000209: protein polyubiquitination
GO:0016604: nuclear body
GO:1904377: positive regulation of protein localization to cell periphery
GO:0035264: multicellular organism growth
GO:0033143: regulation of intracellular steroid hormone receptor signaling pathway
GO:0050840: extracellular matrix binding
GO:0000724: double-strand break repair via homologous recombination
GO:0006974: cellular response to DNA damage stimulus
GO:0009141: nucleoside triphosphate metabolic process
GO:0004177: aminopeptidase activity
GO:0000956: nuclear-transcribed mRNA catabolic process
GO:0035196: production of miRNAs involved in gene silencing by miRNA
GO:0006623: protein targeting to vacuole
GO:0019080: viral gene expression
GO:0002532: production of molecular mediator involved in inflammatory response
GO:0000049: tRNA binding
GO:0005732: sno(s)RNA-containing ribonucleoprotein complex
GO:0031333: negative regulation of protein-containing complex assembly
GO:0047485: protein N-terminus binding
GO:0048812: neuron projection morphogenesis
GO:0030374: nuclear receptor coactivator activity
GO:0008081: phosphoric diester hydrolase activity
GO:0007099: centriole replication
GO:0098732: macromolecule deacylation
GO:0106311: protein threonine kinase activity
GO:1902083: negative regulation of peptidyl-cysteine S-nitrosylation
GO:0003712: transcription coregulator activity
GO:0022604: regulation of cell morphogenesis
GO:0008134: transcription factor binding
GO:0009411: response to UV
GO:0001221: transcription coregulator binding
RI
SE
A5SS
A3SS
MXE

**Supplementary figure 4:** Cluster analysis of enriched pathways of the AS genes in the Erastin treatment group. a: Cluster analysis reveals 2 major clusters, a gene transcription relevant cluster and a translation cluster. b: Analysis of the contribution of different AS types to each node in the clusters confirms the importance of RI in regulating transcription and translation in Erastin. A3SS also appears to play significant role in Erastin.

a

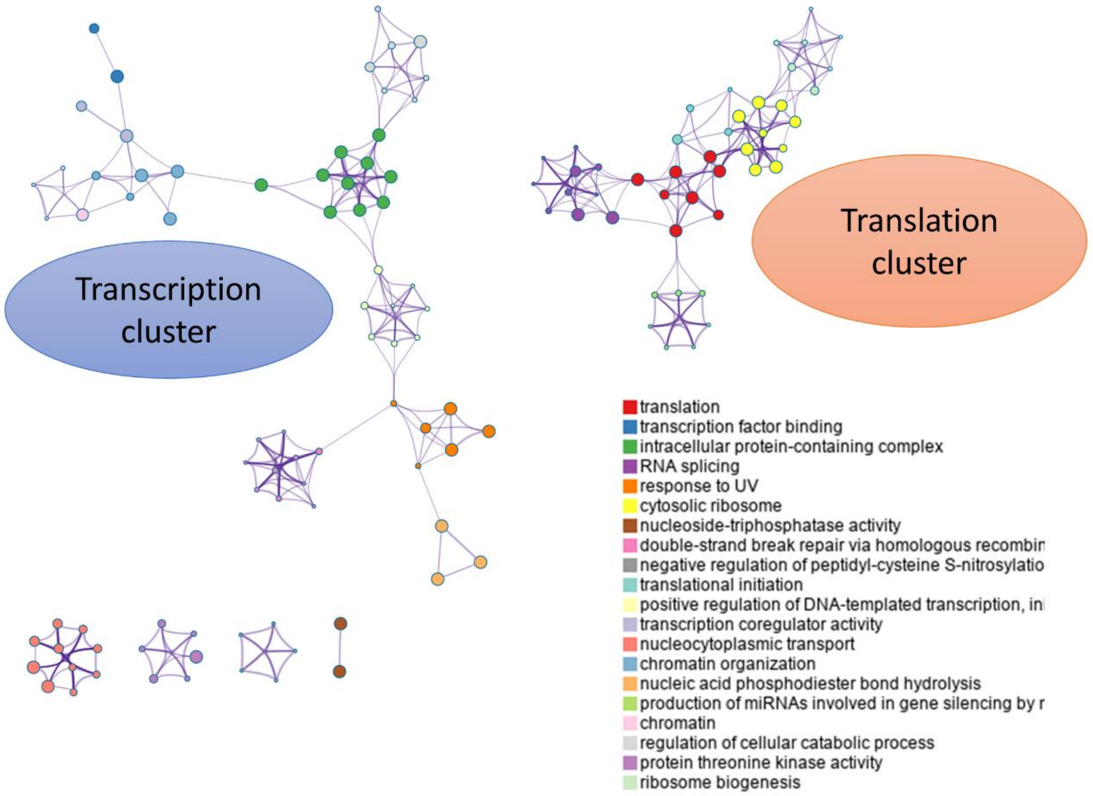

Transcription cluster
Translation cluster
translation
transcription factor binding
intracellular protein-containing complex
RNA splicing
response to UV
cytosolic ribosome
nucleoside-triphosphatase activity
double-strand break repair via homologous recombin
negative regulation of peptidyl-cysteine S-nitrosylatio
translational initiation
positive regulation of DNA-templated transcription, in
transcription coregulator activity
nucleocytoplasmic transport
chromatin organization
nucleic acid phosphodiester bond hydrolysis
production of miRNAs involved in gene silencing by r
chromatin
regulation of cellular catabolic process
protein threonine kinase activity
ribosome biogenesis


b

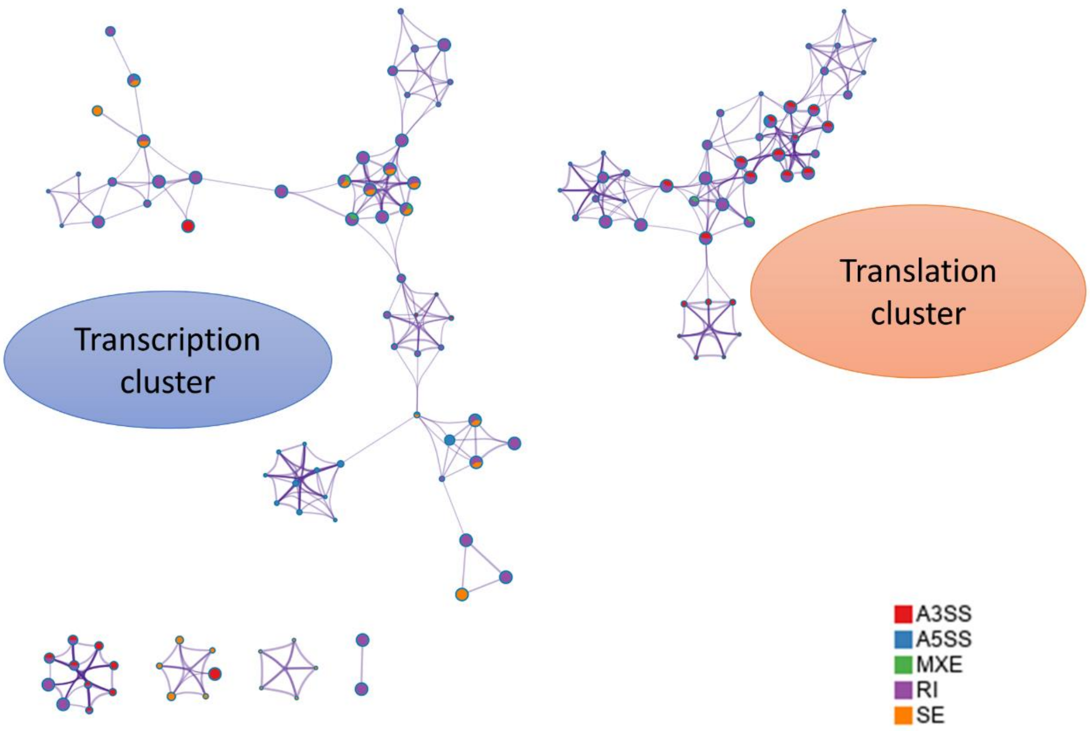

Transcription cluster
Translation cluster
A3SS
A5SS
MXE
RI
SE

**Supplementary figure 5:** Extended GO analysis of the RSL3 AS dataset. Different AS event impact different processes, but the dominance of RI observed in Erastin was not as strong in the RSL3 dataset. (Supplemental to figure 4d).

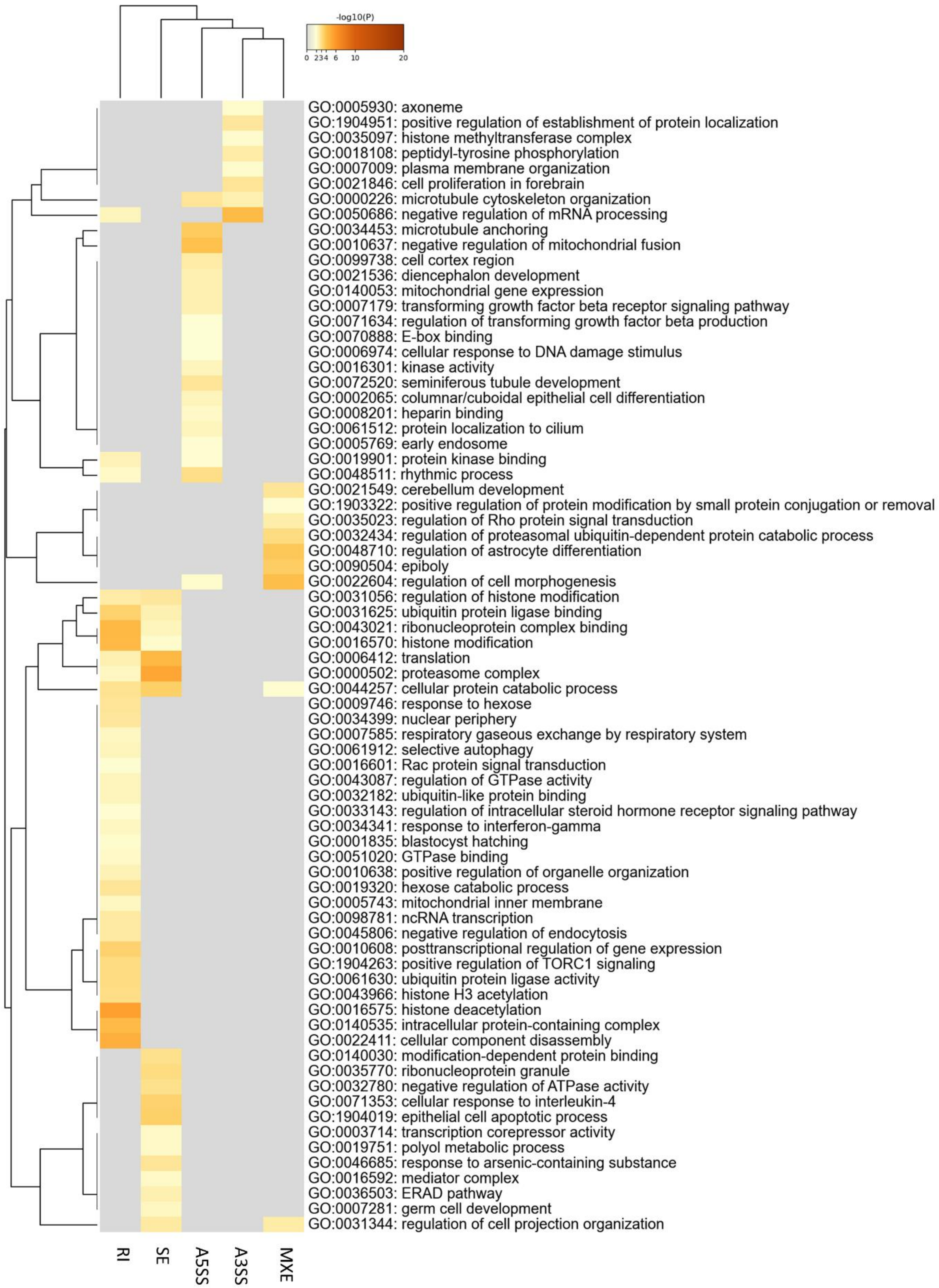
-log10(P)
0
2
3
4
6
10
20
GO:0005930: axoneme
GO:1904951: positive regulation of establishment of protein localization
GO:0035097: histone methyltransferase complex
GO:0018108: peptidyl-tyrosine phosphorylation
GO:0007009: plasma membrane organization
GO:0021846: cell proliferation in forebrain
GO:0000226: microtubule cytoskeleton organization
GO:0050686: negative regulation of mRNA processing
GO:0034453: microtubule anchoring
GO:0010637: negative regulation of mitochondrial fusion
GO:0099738: cell cortex region
GO:0021536: diencephalon development
GO:0140053: mitochondrial gene expression
GO:0007179: transforming growth factor beta receptor signaling pathway
GO:0071634: regulation of transforming growth factor beta production
GO:0070888: E-box binding
GO:0006974: cellular response to DNA damage stimulus
GO:0016301: kinase activity
GO:0072520: seminiferous tubule development
GO:0002065: columnar/cuboidal epithelial cell differentiation
GO:0008201: heparin binding
GO:0061512: protein localization to cilium
GO:0005769: early endosome
GO:0019901: protein kinase binding
GO:0048511: rhythmic process
GO:0021549: cerebellum development
GO:1903322: positive regulation of protein modification by small protein conjugation or removal
GO:0035023: regulation of Rho protein signal transduction
GO:0032434: regulation of proteasomal ubiquitin-dependent protein catabolic process
GO:0048710: regulation of astrocyte differentiation
GO:0090504: epiboly
GO:0022604: regulation of cell morphogenesis
GO:0031056: regulation of histone modification
GO:0031625: ubiquitin protein ligase binding
GO:0043021: ribonucleoprotein complex binding
GO:0016570: histone modification
GO:0006412: translation
GO:0000502: proteasome complex
GO:0044257: cellular protein catabolic process
GO:0009746: response to hexose
GO:0034399: nuclear periphery
GO:0007585: respiratory gaseous exchange by respiratory system
GO:0061912: selective autophagy
GO:0016601: Rac protein signal transduction
GO:0043087: regulation of GTPase activity
GO:0032182: ubiquitin-like protein binding
GO:0033143: regulation of intracellular steroid hormone receptor signaling pathway
GO:0034341: response to interferon-gamma
GO:0001835: blastocyst hatching
GO:0051020: GTPase binding
GO:0010638: positive regulation of organelle organization
GO:0019320: hexose catabolic process
GO:0005743: mitochondrial inner membrane
GO:0098781: ncRNA transcription
GO:0045806: negative regulation of endocytosis
GO:0010608: posttranscriptional regulation of gene expression
GO:1904263: positive regulation of TORC1 signaling
GO:0061630: ubiquitin protein ligase activity
GO:0043966: histone H3 acetylation
GO:0016575: histone deacetylation
GO:0140535: intracellular protein-containing complex
GO:0022411: cellular component disassembly
GO:0140030: modification-dependent protein binding
GO:0035770: ribonucleoprotein granule
GO:0032780: negative regulation of ATPase activity
GO:0071353: cellular response to interleukin-4
GO:1904019: epithelial cell apoptotic process
GO:0003714: transcription corepressor activity
GO:0019751: polyol metabolic process
GO:0046685: response to arsenic-containing substance
GO:0016592: mediator complex
GO:0036503: ERAD pathway
GO:0007281: germ cell development
GO:0031344: regulation of cell projection organization
RI
SE
A5SS
A3SS
MXE

**Supplementary figure 6:** Cluster analysis of the enriched pathways in the AS dataset after RSL3 treatment. No major clusters linked to transcription or translation were observed, contrary to Erastin.

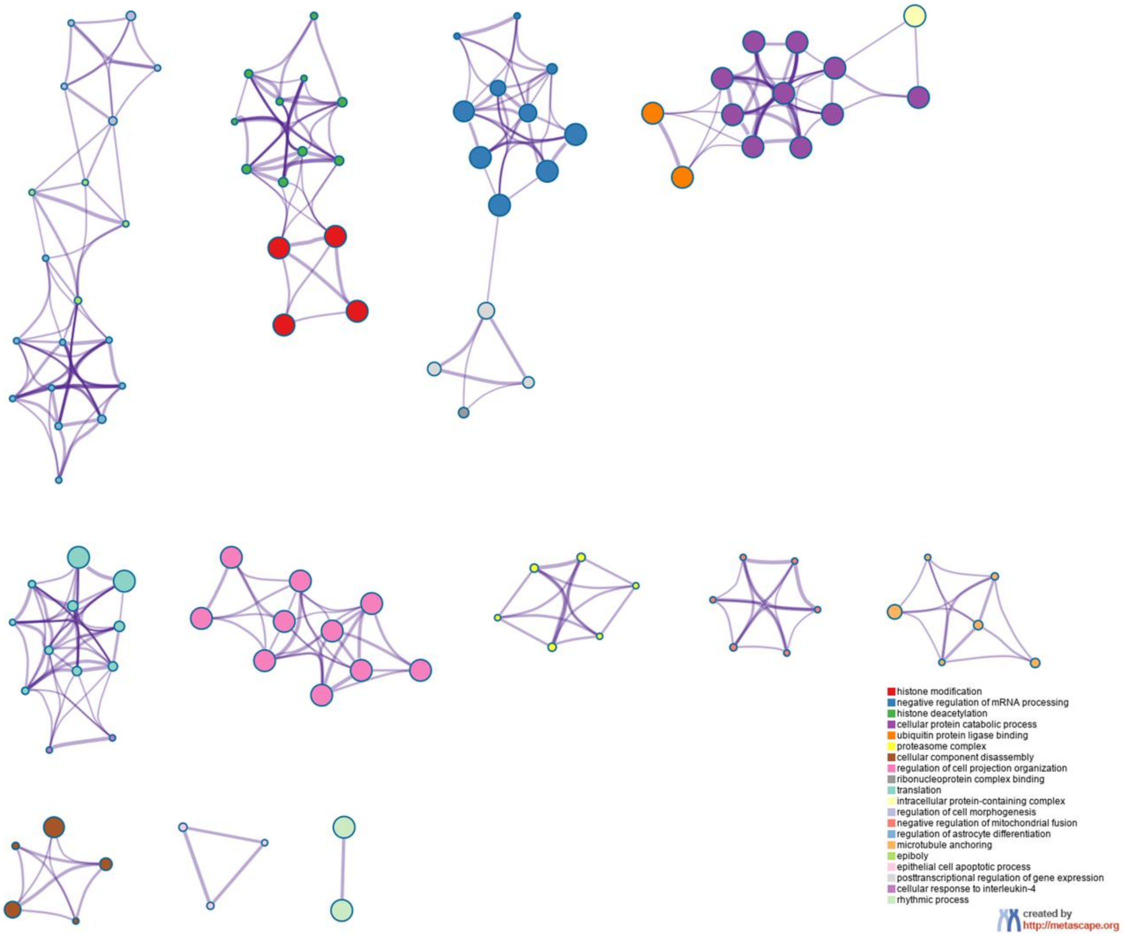

**Supplementary figure 7:** MD plots showing DEGs in the Erastin and RSL3 co-treatment groups with CHX and ActD vs ActD or CHX treatment only groups. No significant patterns were observed in this comparison.

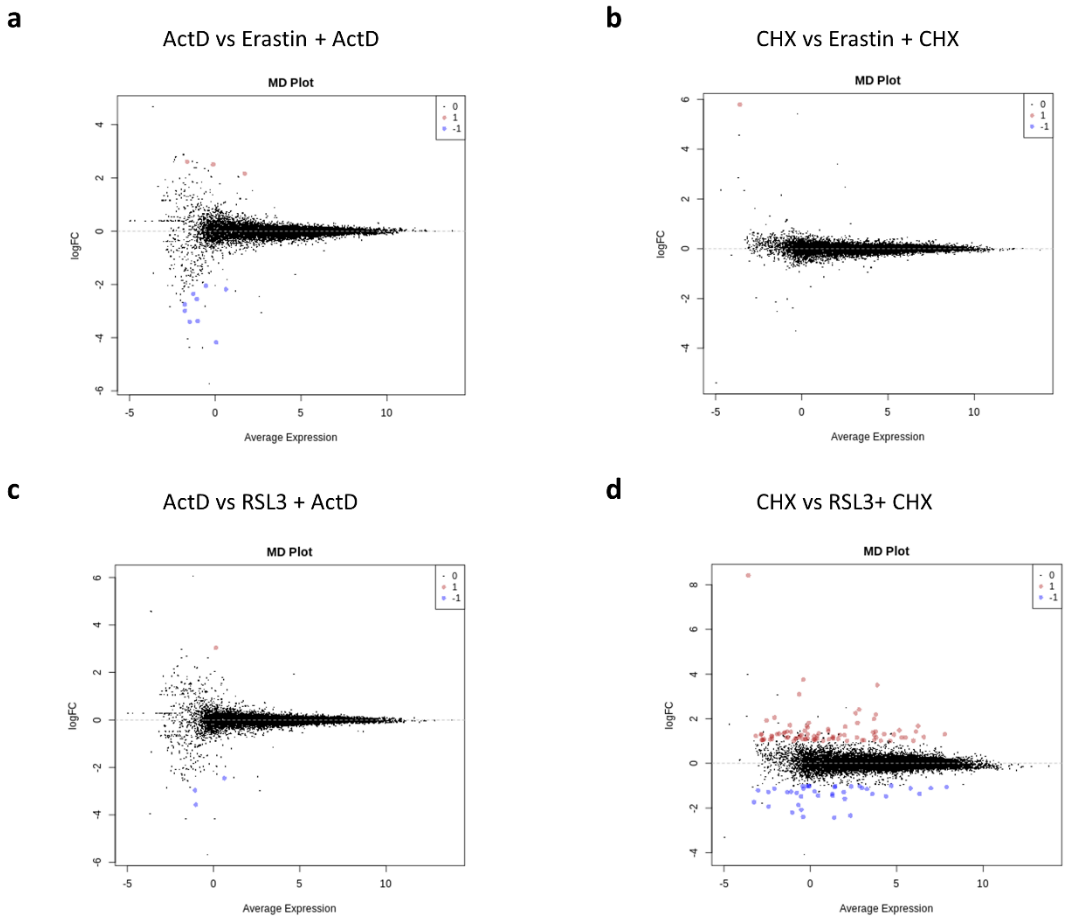

**Supplementary figure 8:** Continued from figure 5c, cluster analysis for nodes that are up or downregulated after CHX treatment and the impact on gene expression and other clusters.

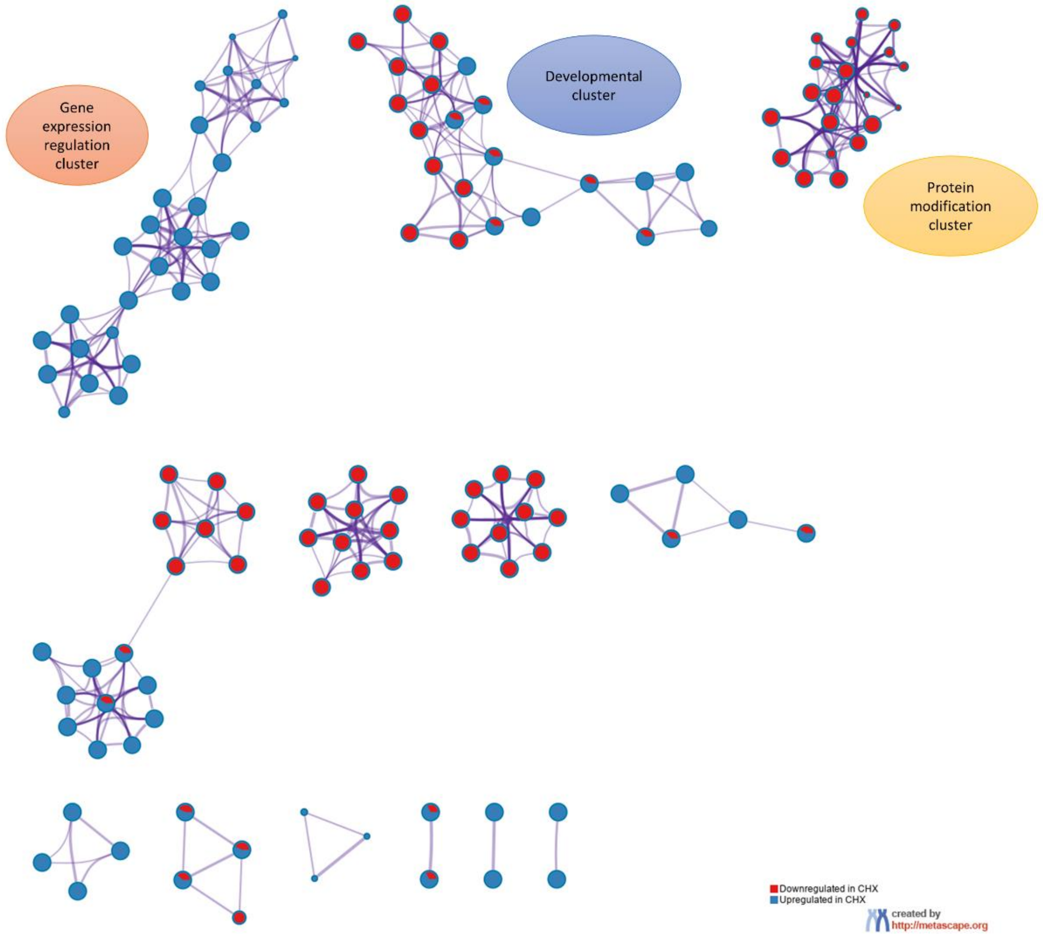

**Supplementary figure 9: mRNA stability analysis in ferroptosis stress response.** a: Methodology for identifying changes in gene stability by using ActD properties. b: GO analysis for the differentially stabilized genes after Erastin or RSL3 treatment. c: GO analysis for the differentially destabilized genes after Erastin or RSL3 treatment.

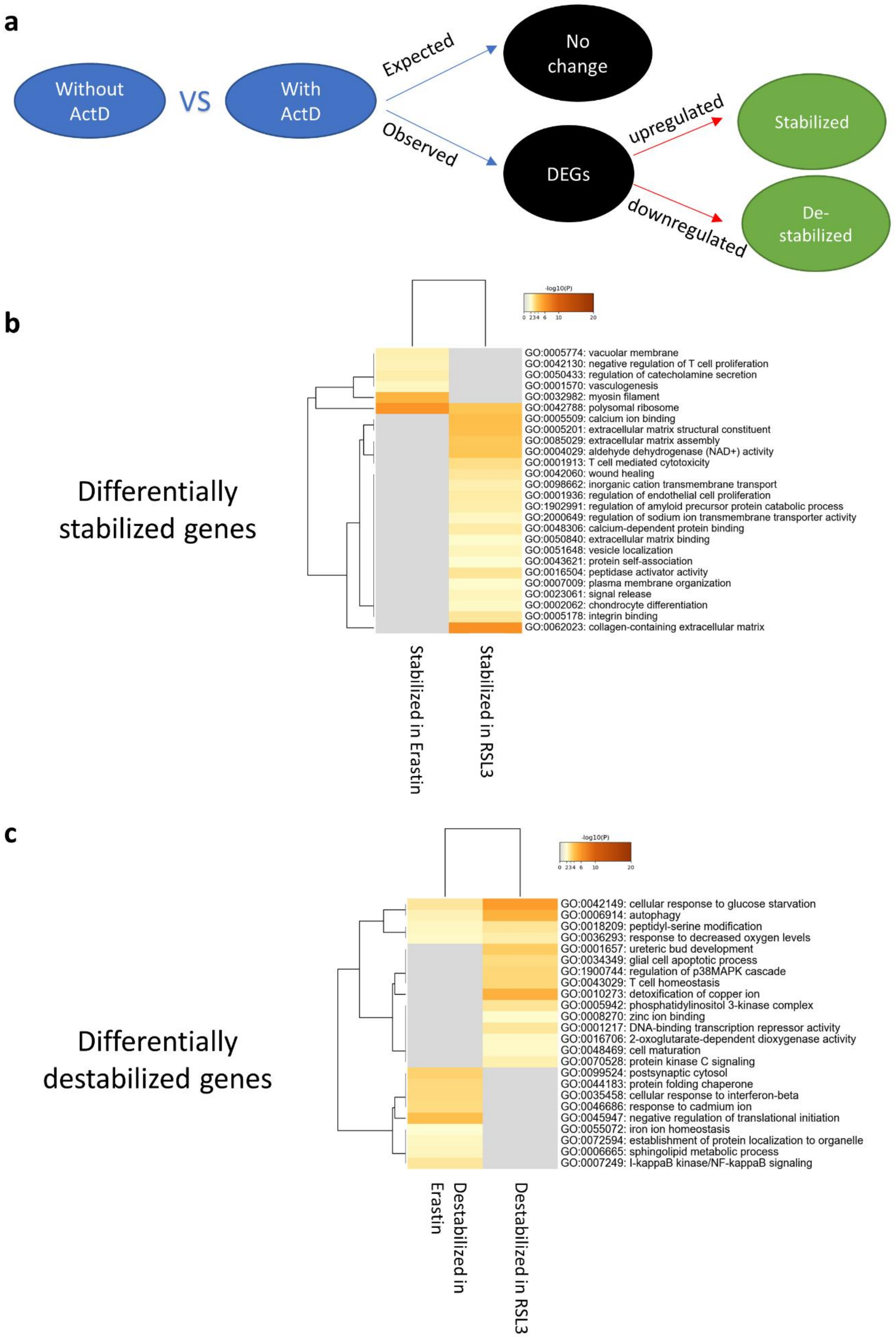
a
Without ActD
VS
With ActD
Expected
No change
Observed
DEGs
upregulated
Stabilized
downregulated
De-stabilized
b
-log10(P)
0 2 3 4 6 10 20
Differentially stabilized genes
GO:0005774: vacuolar membrane
GO:0042130: negative regulation of T cell proliferation
GO:0050433: regulation of catecholamine secretion
GO:0001570: vasculogenesis
GO:0032982: myosin filament
GO:0042788: polysomal ribosome
GO:0005509: calcium ion binding
GO:0005201: extracellular matrix structural constituent
GO:0085029: extracellular matrix assembly
GO:0004029: aldehyde dehydrogenase (NAD+) activity
GO:0001913: T cell mediated cytotoxicity
GO:0042060: wound healing
GO:0098662: inorganic cation transmembrane transport
GO:0001936: regulation of endothelial cell proliferation
GO:1902991: regulation of amyloid precursor protein catabolic process
GO:2000649: regulation of sodium ion transmembrane transporter activity
GO:0048306: calcium-dependent protein binding
GO:0050840: extracellular matrix binding
GO:0051648: vesicle localization
GO:0043621: protein self-association
GO:0016504: peptidase activator activity
GO:0007009: plasma membrane organization
GO:0023061: signal release
GO:0002062: chondrocyte differentiation
GO:0005178: integrin binding
GO:0062023: collagen-containing extracellular matrix
Stabilized in Erastin
Stabilized in RSL3
c
-log10(P)
0 2 3 4 6 10 20
Differentially destabilized genes
GO:0042149: cellular response to glucose starvation
GO:0006914: autophagy
GO:0018209: peptidyl-serine modification
GO:0036293: response to decreased oxygen levels
GO:0001657: ureteric bud development
GO:0034349: glial cell apoptotic process
GO:1900744: regulation of p38MAPK cascade
GO:0043029: T cell homeostasis
GO:0010273: detoxification of copper ion
GO:0005942: phosphatidylinositol 3-kinase complex
GO:0008270: zinc ion binding
GO:0001217: DNA-binding transcription repressor activity
GO:0016706: 2-oxoglutarate-dependent dioxygenase activity
GO:0048469: cell maturation
GO:0070528: protein kinase C signaling
GO:0099524: postsynaptic cytosol
GO:0044183: protein folding chaperone
GO:0035458: cellular response to interferon-beta
GO:0046686: response to cadmium ion
GO:0045947: negative regulation of translational initiation
GO:0055072: iron ion homeostasis
GO:0072594: establishment of protein localization to organelle
GO:0006665: sphingolipid metabolic process
GO:0007249: I-kappaB kinase/NF-kappaB signaling
Destabilized in Erastin
Destabilized in RSL3

**Supplementary figure 10: Analysis of mRNA stability after CHX treatment.** GO analysis for the differentially stabilized (a) or destabilized (b) genes after Erastin or RSL3 treatment and CHX co-treatment.

**a**

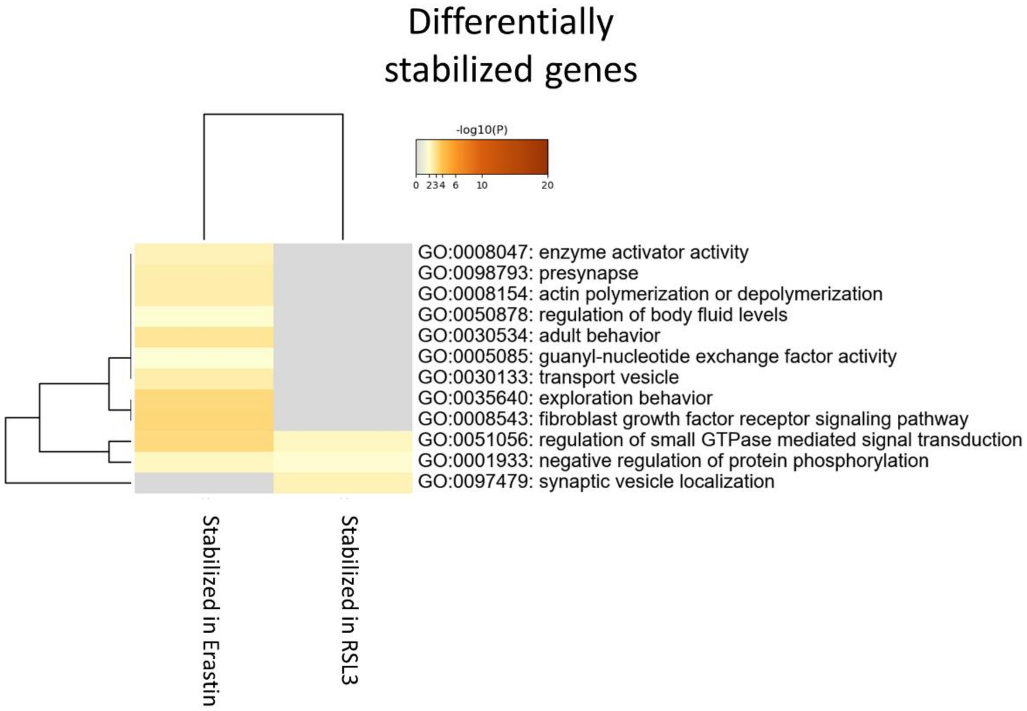
Differentially
stabilized genes
-log10(P)
0 234 6 10 20
GO:0008047: enzyme activator activity
GO:0098793: presynapse
GO:0008154: actin polymerization or depolymerization
GO:0050878: regulation of body fluid levels
GO:0030534: adult behavior
GO:0005085: guanyl-nucleotide exchange factor activity
GO:0030133: transport vesicle
GO:0035640: exploration behavior
GO:0008543: fibroblast growth factor receptor signaling pathway
GO:0051056: regulation of small GTPase mediated signal transduction
GO:0001933: negative regulation of protein phosphorylation
GO:0097479: synaptic vesicle localization
Stabilized in Erastin
Stabilized in RSL3

**b**

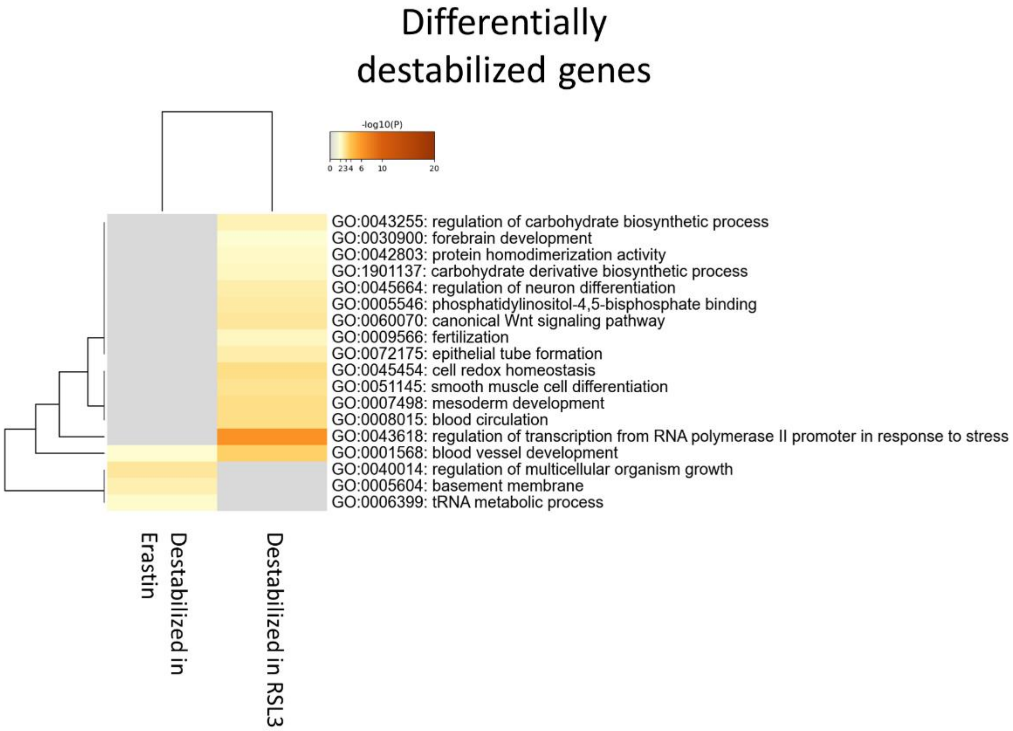
Differentially
destabilized genes
-log10(P)
0 234 6 10 20
GO:0043255: regulation of carbohydrate biosynthetic process
GO:0030900: forebrain development
GO:0042803: protein homodimerization activity
GO:1901137: carbohydrate derivative biosynthetic process
GO:0045664: regulation of neuron differentiation
GO:0005546: phosphatidylinositol-4,5-bisphosphate binding
GO:0060070: canonical Wnt signaling pathway
GO:0009566: fertilization
GO:0072175: epithelial tube formation
GO:0045454: cell redox homeostasis
GO:0051145: smooth muscle cell differentiation
GO:0007498: mesoderm development
GO:0008015: blood circulation
GO:0043618: regulation of transcription from RNA polymerase II promoter in response to stress
GO:0001568: blood vessel development
GO:0040014: regulation of multicellular organism growth
GO:0005604: basement membrane
GO:0006399: tRNA metabolic process
Destabilized in Erastin
Destabilized in RSL3

**Supplementary figure 11:** Scratch test (migration assay) in Mock (a) or Alkbh1+ (b) glioma cells in the presence of Rotenone and Antimycin. (Supplemental to figure 6e)

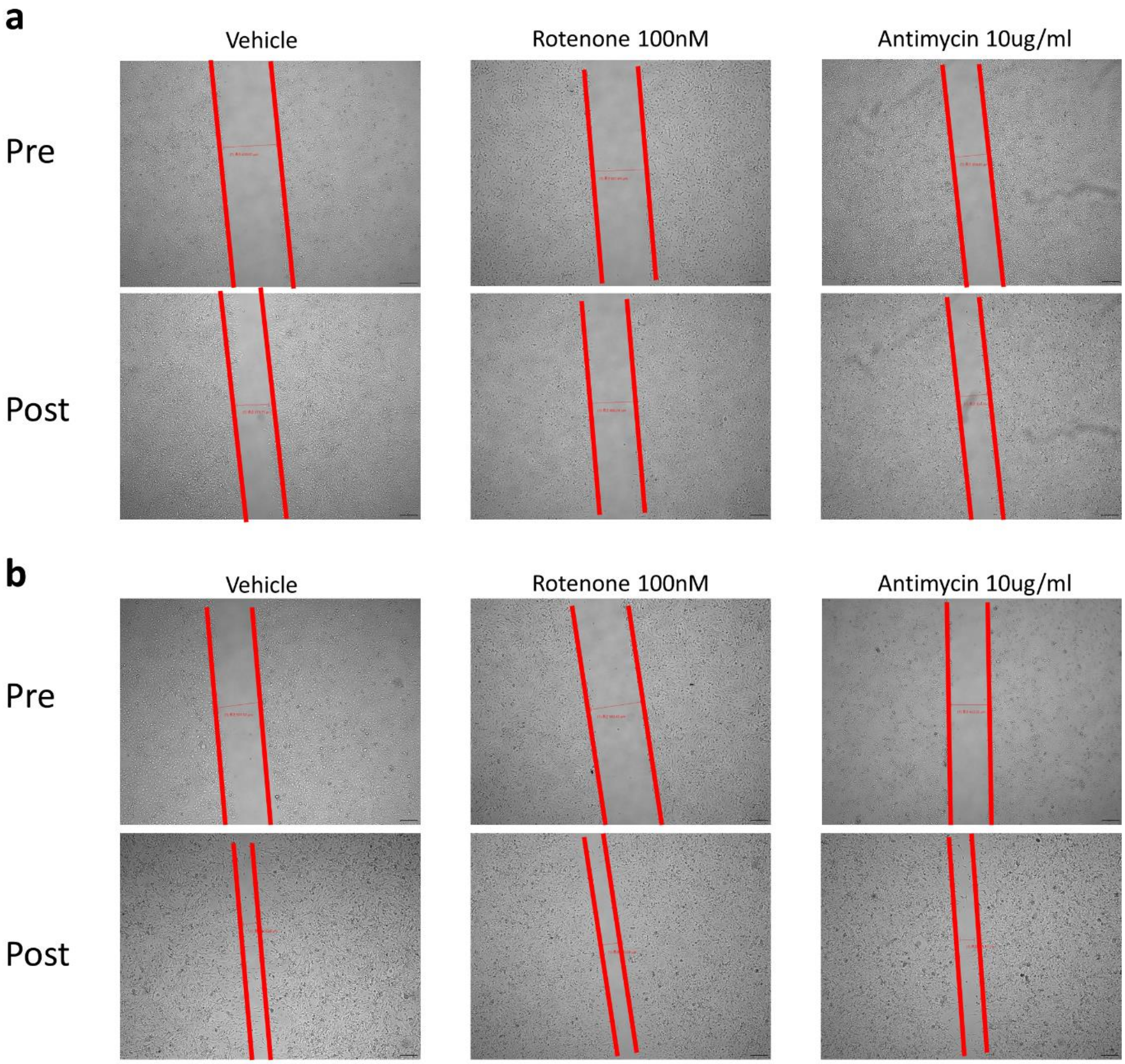
a
Vehicle
Rotenone 100nM
Antimycin 10ug/ml
Pre
Post
b
Vehicle
Rotenone 100nM
Antimycin 10ug/ml
Pre
Post

**Supplementary figure 12:** qPCR gene expression analysis of various genes impacting glioma behavior in Alkbh1+ cells.

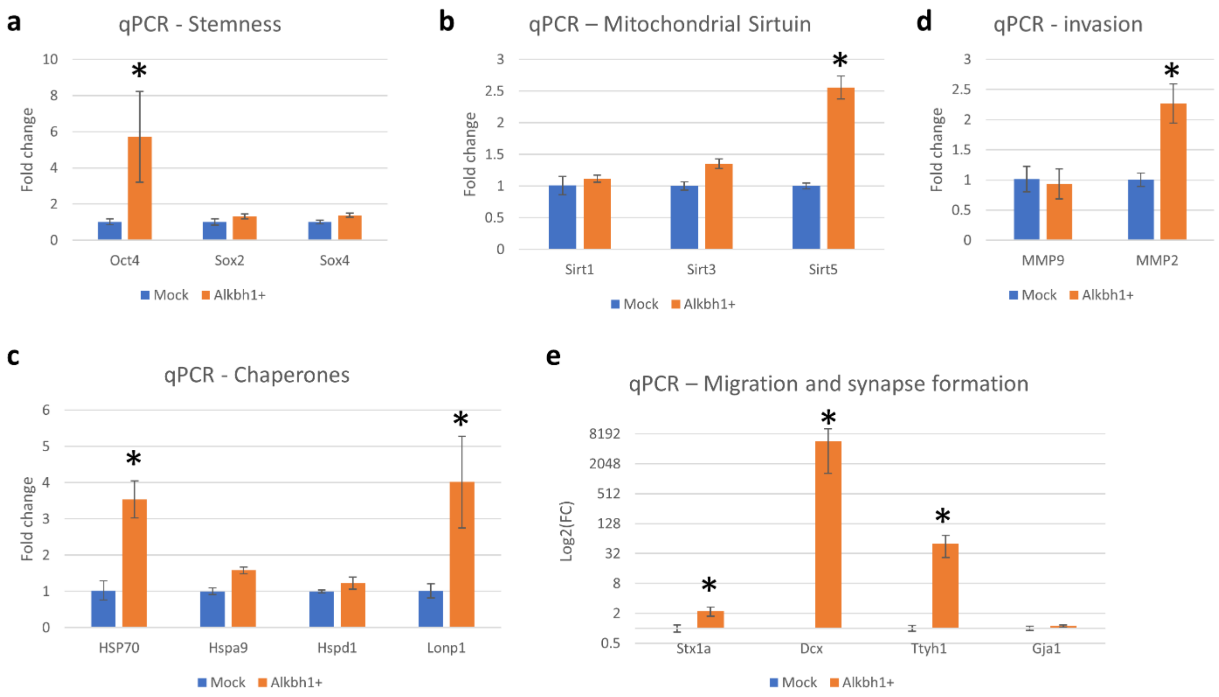

**Supplementary figure 13:** Cluster analysis of enriched pathways in Alkbh1+ cells revealing a DNA and replication cluster that is downregulated and an immune-related cluster that is upregulated. (Supplemental to figure 7b)

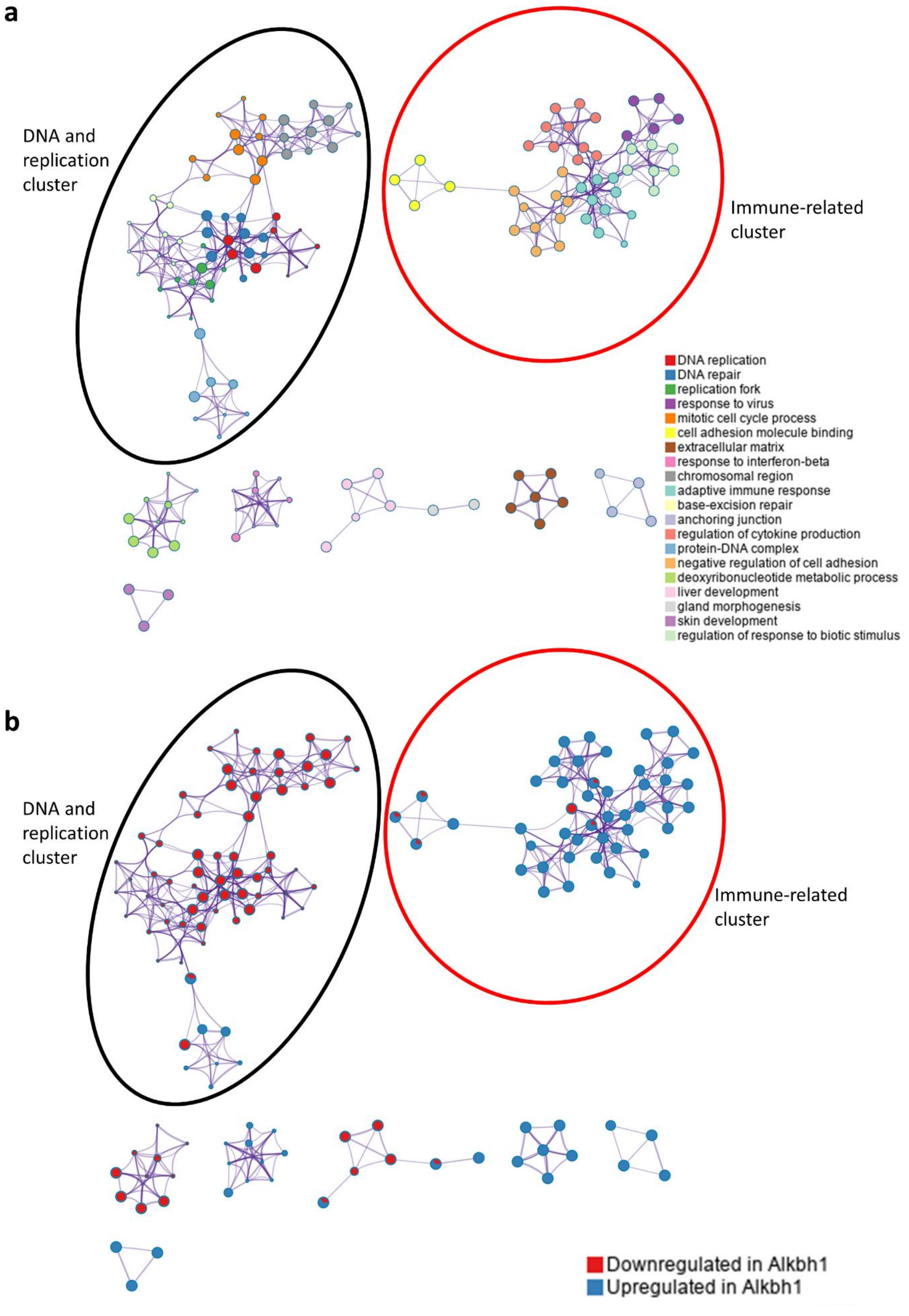
a
DNA and replication cluster
Immune-related cluster
DNA replication
DNA repair
replication fork
response to virus
mitotic cell cycle process
cell adhesion molecule binding
extracellular matrix
response to interferon-beta
chromosomal region
adaptive immune response
base-excision repair
anchoring junction
regulation of cytokine production
protein-DNA complex
negative regulation of cell adhesion
deoxyribonucleotide metabolic process
liver development
gland morphogenesis
skin development
regulation of response to biotic stimulus
b
DNA and replication cluster
Immune-related cluster
Downregulated in Alkbh1
Upregulated in Alkbh1

**Supplementary figure 14: Protein-protein interaction networks in Alkbh1+ cells.** please see supplementary table 3 for MCODE clusters identifications.

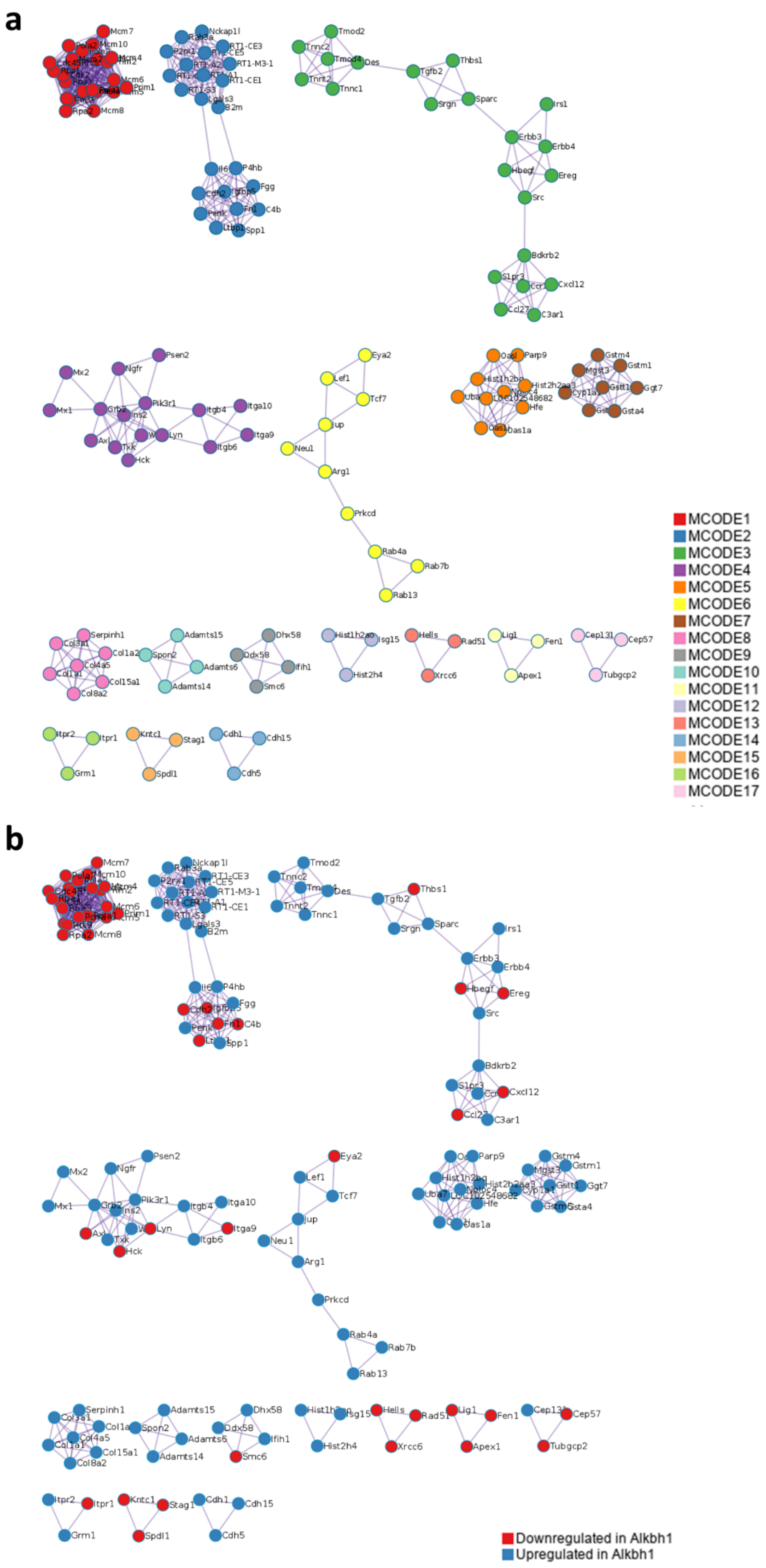
a
MCODE1
MCODE2
MCODE3
MCODE4
MCODE5
MCODE6
MCODE7
MCODE8
MCODE9
MCODE10
MCODE11
MCODE12
MCODE13
MCODE14
MCODE15
MCODE16
MCODE17
b
Downregulated in Alkbh1
Upregulated in Alkbh1
